# Rapid small-scale nanobody-assisted purification of ryanodine receptors for cryo-EM

**DOI:** 10.1101/2024.04.03.587959

**Authors:** Chenyao Li, Katrien Willegems, Tomasz Uchański, Els Pardon, Jan Steyaert, Rouslan G. Efremov

**Affiliations:** Center for Structural Biology, Vlaams Instituut voor Biotechnologie, 1050, Brussels, Belgium; Structural Biology Brussels, Department of Bioengineering Sciences, VIB, 1050, Brussels, Belgium

## Abstract

Ryanodine receptors (RyRs) are large Ca^2+^ release channels residing in the endoplasmic or sarcoplasmic reticulum membrane. Three isoforms of RyRs were identified in mammals, disfunction of which was associated with a series of life-threatening diseases. Advances in structural studies of RyRs are limited by the need for large amounts of native tissue or eukaryotic cell cultures. Here, we report a method that utilizes nanobodies to purify RyRs from only 5 mg of total protein. The purification starting from isolated membranes to cryo-EM grade protein is completed within four hours on the bench and produces protein usable for cryo-EM as we show by solving the structures of rabbit RyR1 and bovine and mouse RyR2 solubilized in detergent, reconstituted into lipid nanodiscs or liposomes. The reported method facilitates structural studies of RyRs directed toward drug development and is useful in the cases where the amount of starting material is limited.

## Introduction

Ryanodine receptors (RyR) are intracellular ion channels that regulate Ca^2+^ flow from the endoplasmic or sarcoplasmic reticulum (ER or SR) into the cytoplasm ^1^. Three isoforms of ryanodine receptor, RyR1, RyR2, and RyR3 were identified in mammals ^2^. RyR1 and RyR2 release Ca^2+^ from SR during contraction of striated and cardiac muscles, respectively ^2^. Hundreds of mutations in RyRs were linked to life threatening diseases. Disfunction of RyR1 causes muscular dystrophy ^3^, malignant hyperthermia (MH) ^4^, congenital myopathies like central core disease (CCD) ^4^, multiminicore disease (MmD) ^5,6^ and atypical periodic paralysis (APP) ^7^. The RyR2 isoform is the major RyR isoform expressed in heart; its disfunction leads to heart failure ^8–10^, catecholaminergic polymorphic ventricular tachycardia (CPVT) ^11^, arrhythmogenic right ventricular dysplasia type 2 (ARVD2) diseases ^12^, Duchenne muscular dystrophy (DMD) ^13^ and dysregulation of insulin release and glucose homeostasis ^14^. RyR3 is an isoform expressed in various organs such as brain, skeletal muscle, liver, and testes ^15–22^. It was implicated in the regulation of blood pressure ^21^, Alzheimer’s disease ^22^, and morphological abnormalities of adrenal gland and hypertrophy of liver ^23^.

RyRs are the largest known ion channels; they form homo tetramers with molecular weight exceeding 2 MDa. A significant part of our mechanistic understanding of RyRs at the molecular level comes from cryo-EM structures of RyR1 and RyR2 isoforms ^24–35^. Each RyR protomer can be divided into 17 structural domains which form a large cytoplasm-exposed crown and a membrane-embedded voltage-gated channel-like domain.

Ca^2+^ release by RyRs is regulated and modulated by interaction with small molecules including Ca^2+^, ATP, caffein and ryanodine ^36^, and interaction with proteins among which are FKBP, calmodulin, and S100A ^37^. Many but not all of them were structurally mapped by cryo-EM and X-ray crystallography ^38^.

Whilst structural data help to understand the effects of mutations and ligand interactions on the gating of RyRs, more structural information is needed for structure-based drug development, including the structures of human RyRs, disease-model RyRs and complexes of RyR with channel modulators. The availability of RyR for structural studies is limited by the availability of native tissues and low levels of protein expression in cell lines ^34,39^. Common purification strategies of solubilized RyRs are ion exchange combined with size-exclusion chromatography, sucrose gradient fractionation, and affinity purification relying on replacing the natively bound FKBP with FKBP carrying an affinity tag ^35,40–42^. Independently of a specific protocol, purification is time consuming and requires a significant amount of starting material to obtain protein amounts suitable for structural analysis. Therefore, protein purification presents a bottleneck for the structural analysis. Further structural studies using cryo-EM, including those that involve the high throughput screening of small molecule binders in the pharmaceutical context would strongly benefit from a rapid small-scale protein purification procedure.

Here, we report a nanobody-based affinity purification method suitable for purifying RyRs solubilized in detergent (RyR1/2-DT), reconstituted into lipid liposomes (RyR1-LP) and into lipid nanodiscs (RyR1-ND). This method requires only 5 mg of total SR membrane protein as starting material. The complete purification takes around 4 h and can be performed on a bench without using an FPLC system. The purified protein can be directly used to prepare cryo-EM grids for structural analysis. We benchmarked the method by solving cryo-EM structures of RyR1 and RyR2 up to a resolution of 3.1 Å in various functional conformational states, using different lipid mimetics and structurally characterized details of the RyR-nanobody interaction.

## Results

### Production and selection of RyR-specific nanobodies

We thought of circumventing the typical long purification procedure of RyRs by developing an affinity-based purification employing nanobodies (Nbs), derivatives of heavy-chain-only camelid antibodies ^43^. As the first step, our goal was to select Nbs with a high affinity for all the RyR isoforms. Because RyR1 is the easiest to purify in amounts sufficient for immunization of llamas, and because mammalian RyR1, RyR2 and RyR3 share around 66% sequence identity, we hypothesized that a significant fraction of nanobodies raised against RyR1 would display a cross-reactivity against RyR2 and RyR3. Therefore, we used rabbit RyR1 reconstituted into amphipols (A8-35) for immunizing the llamas. After initial Nb selection by phage display ^44^, 38 Nbs were screened by ELISA for reactivity against rabbit RyR1 and 25 of them were retained. A subset of 17 Nbs was screened by ELISA against bovine RyR2 and 7 Nbs were retained for further characterization (**Supplementary Figure 1**).

Pull-down assays of rabbit RyR1 and bovine RyR2 from solubilized SR membranes were used to characterize the Nb binding efficiency (**Figure 1**). All selected Nbs bound to both RyR1 and RyR2, however the ratio between the amount of Nb and the amount of bound RyRs measured by the intensity of the SDS-PAGE bands varied between different Nbs. Thus, Nb9657 pulled down the largest quantities of RyR1 and the amount of the pulled down RyR1 remained unchanged upon a 25-fold dilution of the solubilizate. Nb9657 was the best RyR1 binder whereas nanobodies Nb9657 and Nb9662 were the two best RyR2 binders.

**Figure 1.**
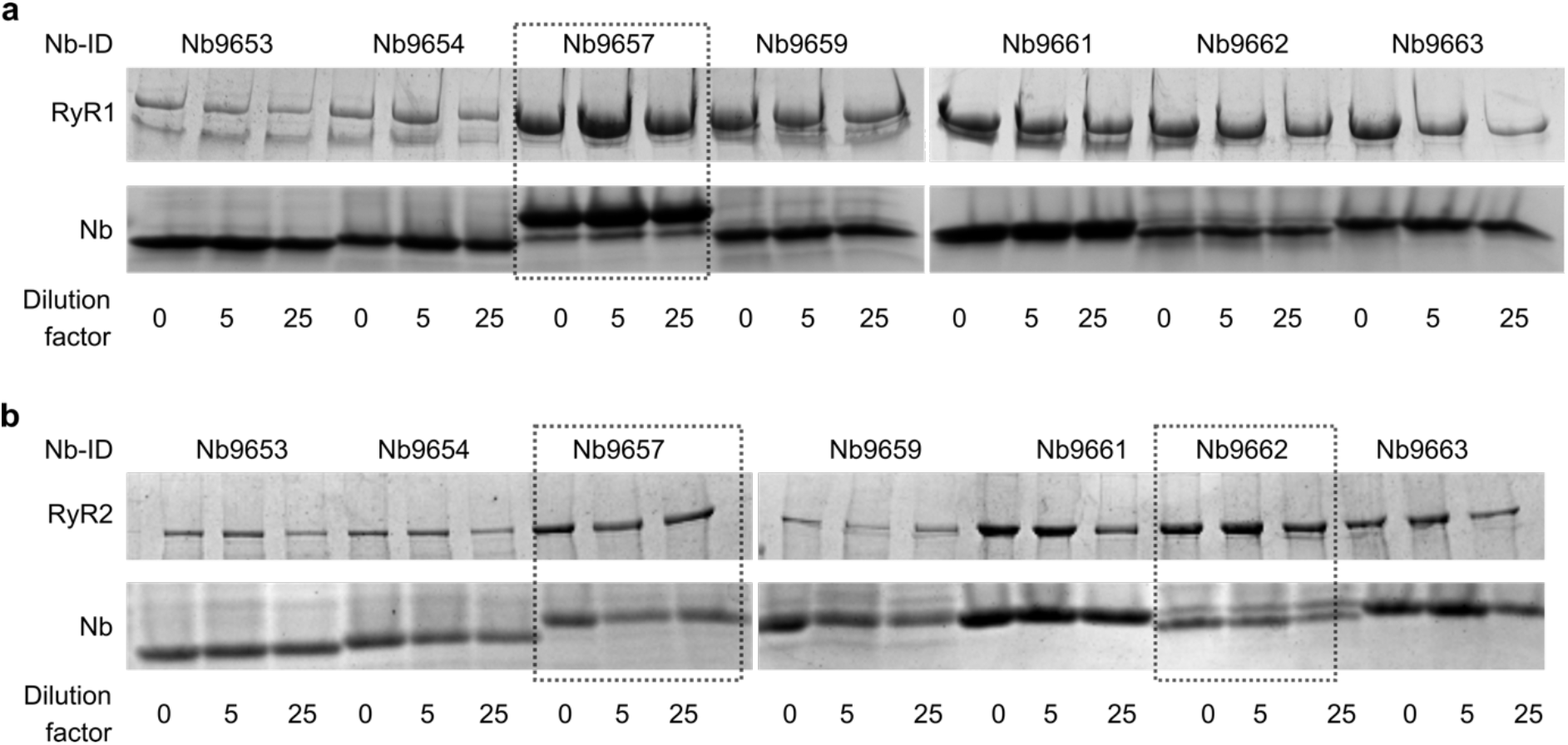
Pull-downs of RyR1 (panel **a**) and RyR2 (panel **b**) with seven selected nanobodies. The nanobodies were coupled to Ni-NTA magnetic beads via His-tag. Coomassie blue stained SDS-PAGEs bands corresponding to RyR monomer (MW ∼ 565kDa) and to Nbs (MW ∼ 15kDa) are shown. The dilution factors are indicated under each lane.

### RyR1 Nb-based purification in lipid nanodiscs

Due to the higher stability and abundance of RyR1, compared to RyR2, the purification and cryo-EM grid preparation was first established for RyR1 and then optimized for RyR2. To provide a native-like lipid environment for RyR1 we developed a purification protocol that included reconstitution of RyR1 into MSP-based lipid nanodiscs^45^ The resulting purification strategy consisted of trapping RyR1 using the Nb9657 immobilized on Ni-NTA magnetic beads followed by co-elution of Nbs and bound RyR1 with imidazole (**Figure 2a, b**). Because the elution buffer contained lipids, required for stabilizing the native state of RyR1, the reconstitution into the nanodiscs was accomplished by supplementing eluted protein with MSP1E3D1^46^ followed by slow dilution. The MSP:lipid ratio for the reconstitution was optimized empirically and the optimal value was found to be identical to the theoretical value of 1 MSP per 130 lipids. Finally, 0.2% of fOM (fluorinated octyl maltoside) was added to reduce aggregation of the reconstituted RyR1. The excess of empty lipid nanodiscs was removed by repeating the nanobody affinity purification step in a buffer with CHAPS replaced with fOM. Additionally, this purification step improved the purity of RyR1 (**Figure 2b**). Finally, the excess of co-eluted Nb9657 and imidazole were removed using a spin column packed with a size-exclusion resin (**Figure 2b** lane S1).

**Figure 2.**
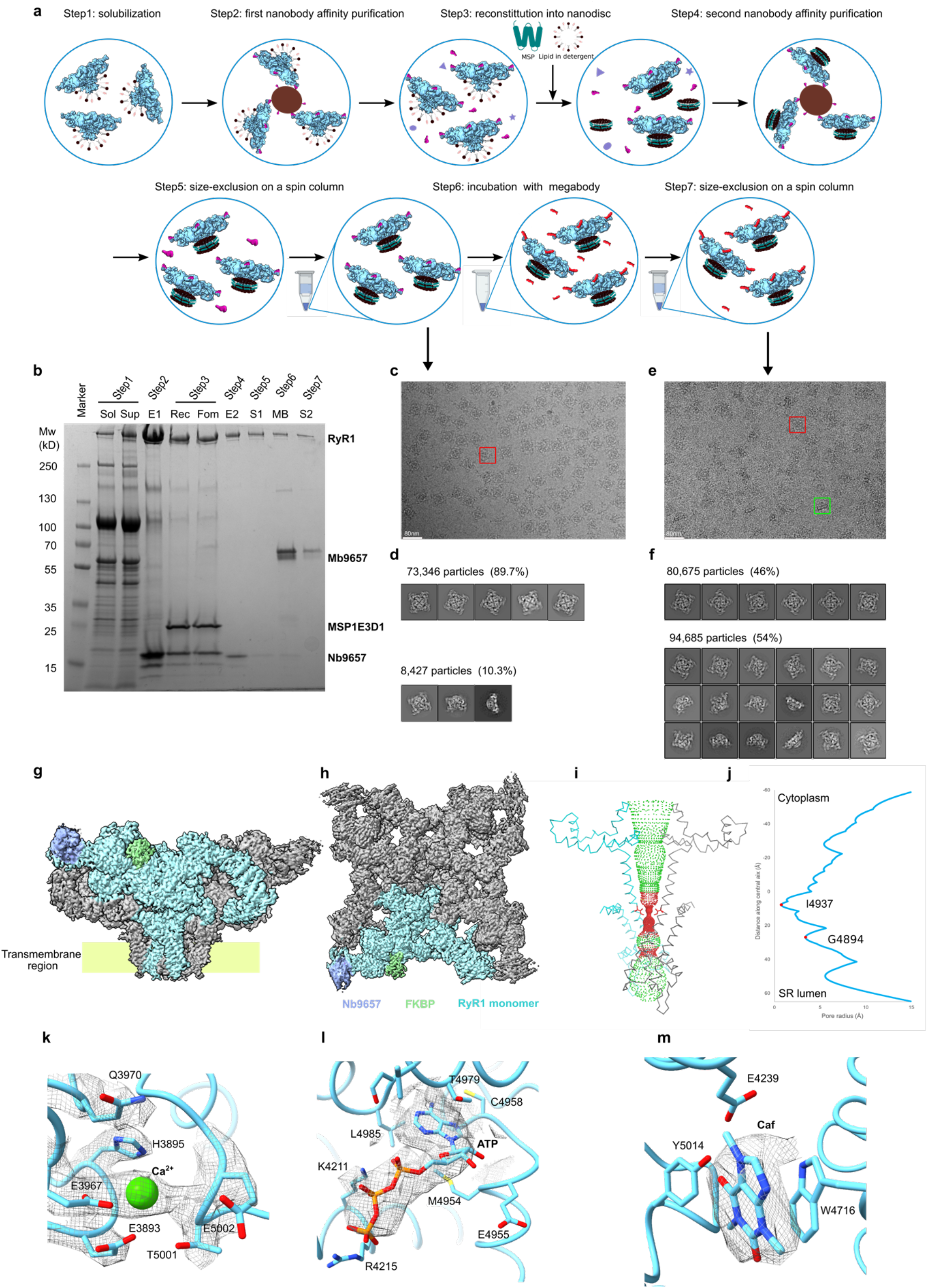
Nb-based purification and single particle cryo-EM of RyR1-ND. **a**, Schematic diagram of nanobody-based affinity purification and reconstitution into lipid nanodiscs. **b,** Coomassie-stained SDS-PAGE during sequential steps of the purification procedure shown in panel **a.** Following abbreviations for lane tags were used: Sol – solubilized SR membranes, Sup - supernatant of ultra-centrifuged Sol, E1 - eluate of the first purification round, Rec - RyR1 reconstituted into lipid nanodisc, Fom - Rec solution with added fOM, E2 - eluate of the second purification round, S1 – eluate from the first spin column, MB - incubation with Mb9657, S2 – eluate from second spin column. **c-f**, Cryo-EM micrographs and 2D classes of RyR1-ND bound to Nb9657 (**c, d**) and to Mb9657 (**e, f**). The red square indicates a top and the green square indicates a side view of RyR1 particle. **g, h,** Top and side view of the cryo-EM map of RyR1-ND complexed with Mb9657. Nb9657, FKBP and RyR1 monomer are colored in purple, green and cyan respectively. **i,** Side view of the pore region of the RyR1-ND structure, two protomers are shown in cyan and gray, and the gate residue I4937 is in red. The channel solvent accessible surface was calculated with HOLE ^48^.The channel regions with radius above 4 Å are shown in green and below 4 Å in red. **j,** Graph of the pore radius shown in **i.** The location of residues I4937 and G4984 are indicated with red dots. **k-m,** density of Ca^2+^, ATP and caffeine in the RyR1-ND map, respectively.

RyR1 eluted at a concentration of around 1 mg/ml and was directly used for the preparation of cryo-EM samples on graphene oxide-coated grids (**Figure 2c**). However, a strong preferential particle orientation with prevailing top views was observed such that the tilted and side views accounted for only 10% of the particles (**Figure 2d**). The lack of side views limited the resolution of the cryo-EM maps to around 4.7 Å (**Supplementary Figure 2**). To overcome the preferred orientation, we used megabodies (Mbs) which, in addition to being commonly used as fiducials, were reported to improve particle orientation distribution ^47^. To this end, we designed a Mb in which Nb9657 was grafted onto the cHopQ scaffold ^47^ (further referred to as Mb9657).

RyR-bound Nb9657 was replaced with Mb9657 by incubating the purified RyR1 with the Mb followed by a second round of spin desalting column to remove the excess of Mb (**Figure 2a, b**). This resulted in an improved orientation distribution of RyR1 particles where around 50% of the particles presented a tilted or side view (**Figure 2e, f**). This allowed obtaining a cryo-EM 3D map of RyR1-Mb9657 complex at the resolution of 3.1 Å (**Figure 2g, h, Supplementary Figures 3, 7** and **Supplementary Table 1**). The resulting map clearly resolved side chains throughout the structure and density for several lipid molecules. The structure was solved in the presence of activating ligands including Ca^2+^, ATP and caffein, for which a clear density was observed in the reconstruction (**Figure 2k, l, m**).

The conformation of RyR1-ND corresponded to the primed state in which Ca^2+^, ATP and caffein are bound whereas the ion channel gate remains closed (**Figure 2i, j**). When compared to the RyR1 primed state structure solved in liposomes (7M6A) ^25^ under similar conditions, the Cα RMSD for the pore and C-terminal domain (residues 4820-5037) was 0.8 Å whereas the global conformation was somewhat different with the crown regions shifted towards the nanodisc with the amplitude at the periphery of around 3 Å (**Supplementary Figure 4f**).

The density of endogenous FBKP was well resolved in the reconstruction (**Figure 2g, h**), which indicates that the purification method is soft enough to co-purify some of the interaction partners. An additional density corresponding to the bound nanobody was observed next to the Repeat 12 domain (**Figure 2g, h**).

The amounts of the reconstituted RyR1 sufficient for structural analysis were purified starting from isolated SR membranes with the total protein content of around 5 mg. The complete purification and reconstitution into the lipid nanodiscs were conducted on a bench and took under 4 hours.

As the next step, we explored the compatibility of different lipid mimetics with the established purification strategy.

### Nb-based purification and cryo-EM of detergent-solubilized RyR1

The nanobody-based affinity purification was also applicable for purifying and solving the structure of the CHAPS-solubilized RyR1 (RyR1-DT). Solubilized RyR1 suitable for structure determination by cryo-EM was obtained following two cycles of trapping RyR1 on Nb-loaded Ni-NTA magnetic beads after which the buffer was exchanged twice on spin columns to remove imidazole and to replace the Nb with Mb (**Figure 3a, b**). To maintain protein stability, the purification was performed in a buffer containing CHAPS (0.8% w/v) and POPC (0.2% w/v), concentration of both of which was reduced during the last size-exclusion step to 0.2% and 0.001% (w/v), respectively, to improve the contrast on cryo-EM grids.

**Figure 3.**
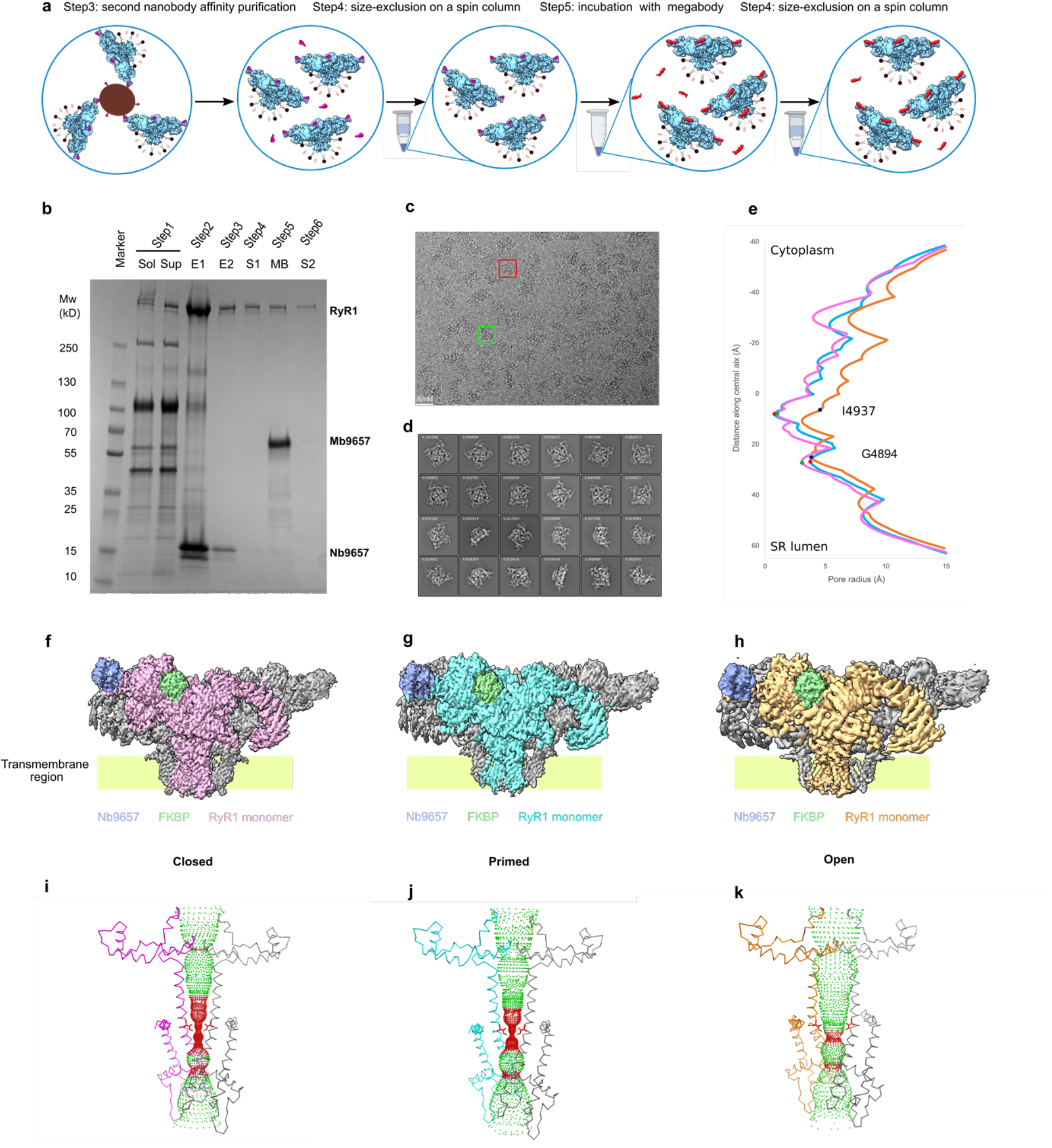
Purification of RyR1-DT. **a,** Schematic diagram of nanobody-based affinity purification of RyR1-DT. Purification steps 1 and 2 were the same as in Figure 2a**. b,** Coomassie-stained SDS-PAGE of the individual steps of RyR1 purification. The lanes are labeled as in Figure 2b. **c-d,** A cryo-EM micrograph and 2D classes of RyR1 in detergent micelles, red square indicates a top view and green square a side view particle. **e,** Plots of the pore radius for closed, primed and open states are shown in pink and cyan and orange, respectively. The positions of I4937 and G4894 are indicated by dots. **f, g, h** Cryo-EM map of RyR1-DT in closed, primed and open states. Monomers of each state are colored in pink, cyan and yellow, nanobody and FKBP are colored purple and green. **I-k,** Side view of pore region of RyR1-DT structure in closed, primed and open state. The solvent accessible volume calculated in HOLE is shown as dotted surface and colored as in Figure 2i.

Purified RyR1-DT was plunge-frozen on graphene-oxide-coated cryo-EM grids in the presence of EGTA. After collecting single-particle datasets (**Figure 3c-d, Supplementary Table 1**) the structure was solved to a resolution of 3.2 Å (**Figure 3f, Supplementary Figures 3 and 9**). The resulting map corresponded to a closed state conformation (**Figure 3e, i**) and closely matched previously determined structure of the CHAPS solubilized rabbit RyR1 in closed state (PDB code 8SEN ^49^, RMSD 0.8 Å for 218 Cα atoms of the pore region residues 4820-5037, the overall RMDS for 4112 Cα atoms was 4.5 Å due to the shift of BSol domain towards the membrane, **Supplementary Figure 4a**).

To verify that the purified RyR1-Mb9657 complex can be stabilized in the open conformation, channel activators including Ca^2+^, ATP and caffein were added to RyR1-DT during the spin column steps and the cryo-EM data were collected from the purified protein. After image processing and 3D classification, the structures of RyR1 in the primed and open conformations were reconstructed to the resolution of 3.3 Å and 4.2 Å (**Figure 3g, h, Supplementary Table 1, Supplementary Figure 8**) to which 84% and 16% of particles contributed, respectively.

Similar to the RyR1-ND, the cryo-EM maps of RyR1-DT also showed density for Nb9657 and native FKBP12. The density of Ca^2+^, ATP and caffeine were observed in the same locations as in the RyR1-ND reconstruction. The primed state structure has a constricted pore, with the gate diameter of less than 1 Å formed by I4937 (**Figure 3e, g, j**), whereas the open state structure has a dilated channel which allows for the passage of Ca^2+^ ions (**Figure 3e, h, k**).

### Structure of Nb-purified RyR1 reconstituted into lipid liposomes

By serendipity, we observed that when the buffer exchange step was performed with CHAPS concentration below CMC (∼ 0.5% ^50^), a considerable fraction of RyR1 particles was reconstituted into lipid liposomes. We have used this to our advantage, and following optimization of the CHAPS concentration, cryo-EM samples containing liposomes with a diameter in the range between 30 and 50 nm were reproducibly generated (**Figure 4**). In the sample, the reconstituted RyR1 particles were observed along with non-reconstituted particles similar to a more sophisticated preparation protocol reported by Melville et. al.^51^ Due to the large size of the cytoplasmic domain of RyR, its structure can be relatively easily reconstructed for the particles reconstituted into liposomes ^51^. It is important to note, that the reconstituted RyR1 particles were observed as side views and did not require Nb to Mb exchange to improve particle orientation distribution. After collecting a cryo-EM dataset and performing 2D classification, we selected classes displaying strong density for the lipid bilayer around the membrane-embedded domain of RyR1 (**Figure 4d**) and reconstructed the structures of RyR1 in the liposomes (RyR-LP) to resolution between 4.5-4.7 Å (**Figure 4e, f, Supplementary Figures 3, 10, Supplementary Table 1**). The pore region had the highest local resolution of 4.1 Å. At lower threshold, the density of bent lipid bilayer was visualized (**Figure 4e, f**), as was expected for the reconstructions from the liposomes.

**Figure 4.**
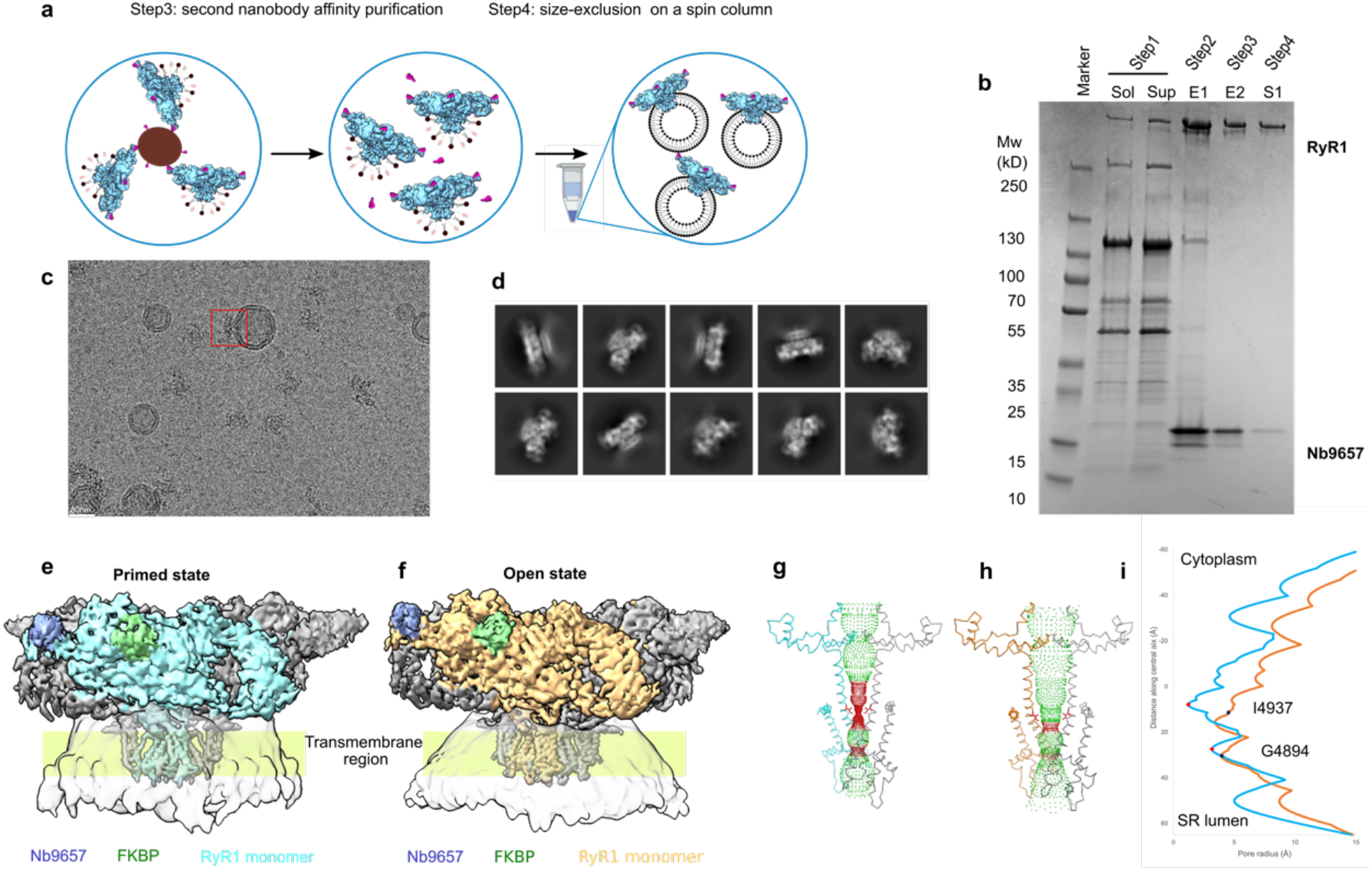
Purification and reconstitution of RyR1 into liposomes and cryo-EM of the reconstituted protein. **a,** Schematic diagram of nanobody-based affinity purification of RyR1-LP. The purification steps 1 and 2 were the same as in Figure 2a**. b,** Coomassie-stained SDS-PAGE of the RyR1 sample at sequential steps of the purification procedure; the lanes labels are the same as in Figure 2b**. c-d,** A cryo-EM micrograph and 2D classes of RyR1-LP reconstituted into liposomes. The 2D classes show particles used for RyR1 map construction. Only classes displaying well-defined density of liposomes were selected. **e-f,** Cryo-EM maps of RyR1-LP in primed and open states, the monomers of the primed and open state are colored in cyan and orange respectively, nanobody and FKBP are colored in purple and green, respectively. **g-h,** Side view of the pore region of RyR1-LP structure in primed and open states. The HOLE^48^ calculated dotted accessible surface and hydrophobic gate residue I4937 are colored the same way in Figure 2i**. i,** Plot of the pore radius is shown for the primed (cyan) and open (orange) states.

The RyR1-LP was prepared in the presence of channel activators (Ca^2+^, ATP and caffeine), consequently the cryo-EM maps of the primed and open states were reconstructed with occupancies of 38 and 62%, respectively (**Figure 4e, f, Supplementary Table 1**). Here the fraction of the open state is higher (62 vs 40%) than was previously reported for RyR1 reconstituted into lipid liposomes ^33^.

This example demonstrates that our purification approach is also suitable to study RyR in a compartmentalized system in which conditions for cytoplasm and lumen-exposed domains may be set differently, even though further optimization will be needed to increase the number of reconstituted particles and improve the resolution of the RyR1 reconstructions.

### Application of Nb-based purification for structural studies of RyR2

Our objective was to establish a purification approach that works for various RyR isoforms, therefore as the next step we applied the purification method to RyR2. RyR2 is the prevalent RyR isoform in the ventricular heart tissue. When compared to striated muscles, the content of RyR2 in the SR membranes extracted from heart tissue is notoriously lower than that of RyR1 extracted from the striated muscles (compare **Figures 2b and 5a**). The lower amount combined with lower stability of RyR2, makes the protein a considerably more challenging target for structural studies. This is reflected in a lower number of RyR2 structures deposited to PDB as compared to RyR1 (47 versus 72, respectively).

**Figure 5.**
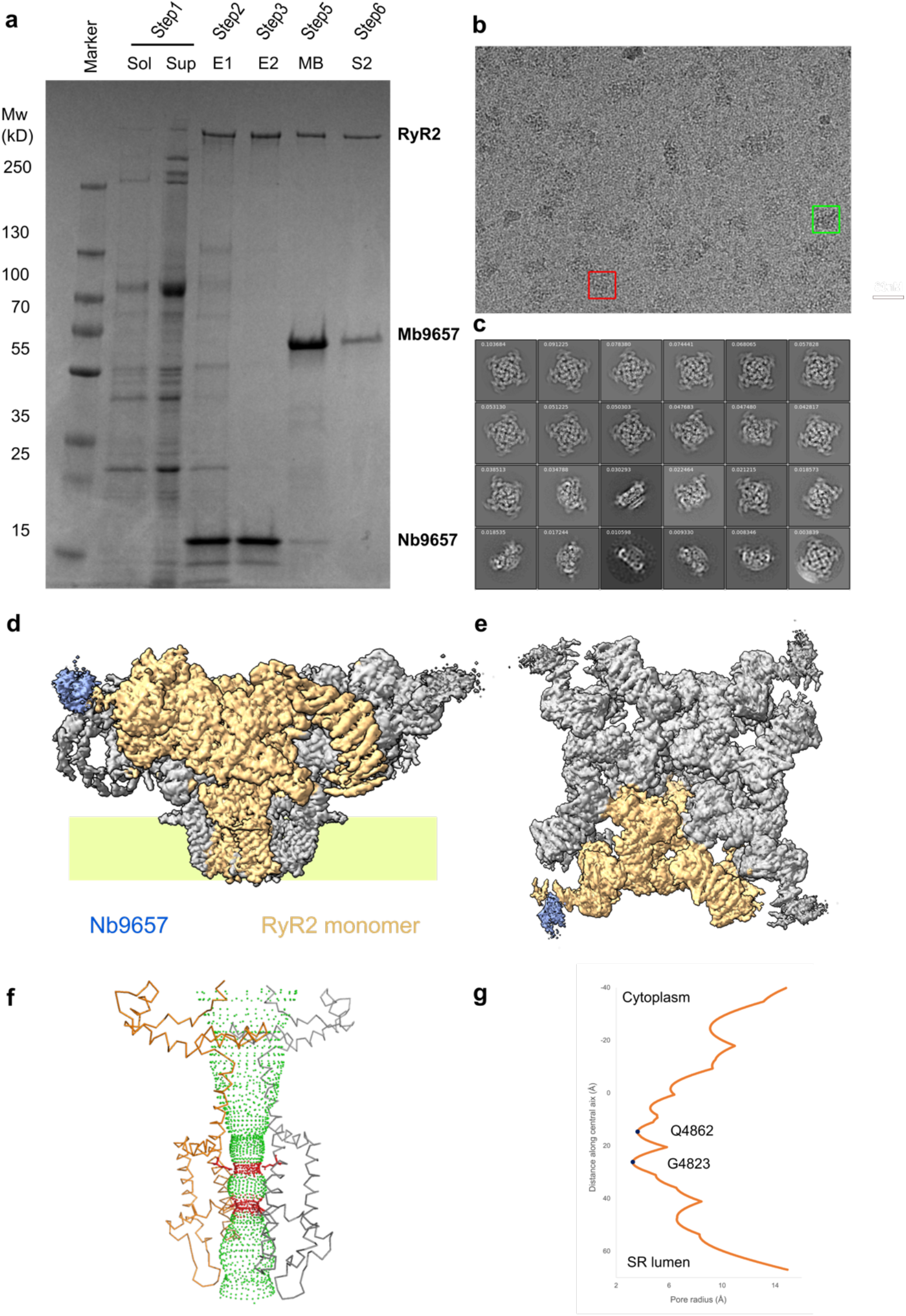
Purification of RyR2-DT. **a,** Coomassie-stained SDS-PAGE of samples taken from the sequential steps of mouse RyR2purification. The lanes are labeled as in Figure 2b**. b-c,** A cryo-EM micrograph and 2D class averages of RyR2-DT, the red and green squares indicate top and side views of the receptor. The 2D classes show particles used for RyR2 map reconstruction. **d-e,** Side and top view of RyR2 3D map, nanobody and a RyR2 monomer are colored in purple and orange, respectively. **f,** Side view of pore region of RyR2-DT structure is shown along with the pore surface calculated with HOLE. The hydrophobic gate residue I4937 and dotted channel surface are colored the same way in Figure 2h**. g,** Plot of the pore radius.

We initially purified RyR2 from cow hearts in lipid nanodiscs and obtained its cryo-EM structure in complex with Nb9662 to 7.5 Å resolution in presence of 10 µM free Ca^2+^ (**Supplementary Figure 5**). The structure was the most consistent with the primed state. Interestingly, in complex with Nb9662, bovine RyR2 formed dimers of RyR2 tetramers in which the tetramer-tetramer contacts were mediated by the nanobodies (**Supplementary Figure 5d**). These dimers disassembled when EDTA was added, suggesting that the Nb-Nb interaction was mediated by divalent ions. Due to the limited supply of cow hearts, we continued the work with naive C57BL6 mouse hearts and changed Nb9662 to Nb9657 which was more stable and was less prone to forming the RyR2 dimers.

The CHAPS-solubilized mouse RyR2 was purified following the same steps as used for RyR1-DT purification (**Figures 2a, b and 5a**). Lower yield and stability of RyR2 required a few modifications of the purification buffers: 1) 20 mM MOPS used for RyR1 was replaced with 50 mM HEPES, 2) a higher concentration of TCEP (5 mM) was used during all purification steps, and 3) the detergent and lipid concentrations were adjusted to 0.6 % CHAPS and 0.3 % POPC. Due to the lower RyR2 content in cardiac SR membranes, an amount of SR membranes with total protein content of around 14 mg was used as starting material. This quantity of membranes was extracted from around 15 mouse hearts.

A single purification yielded 50 µl of pure CHAPS-solubilized RyR2 bound to Mb9657 at a concentration of around 1 mg/ml. This protein was directly used for plunge-freezing on graphene-oxide coated cryo-EM grids and produced grids with well-distributed particles and sufficient particle density for structure determination (**Figure 5b, c**). The RyR2 agonists: Ca^2+^, caffeine and ATP were added during protein purification.

We obtained a cryo-EM map of mouse RyR2-DT at a resolution of 3.4 Å (**Figure 5, Supplementary Figures 3, 11, Supplementary Table 1**). The reconstruction represented the open state of the receptor, and no primed state was detected after 3D classification suggesting that occupancy of the open state was close to 100%. Unlike in the case of RyR1, the density for the endogenous FKBP was missing. This is consistent with lower affinity of FKBP12.6, mainly present in cardiac cells, for RyR2 as compared to the affinity of FKBP12 for RyR1 ^37^. The density for Nb9657 was resolved in the position and orientation identical to that in the RyR1 structure (**Figure 5d, e**).

This mouse RyR2 reconstruction is, to the best of our knowledge, the first reported RyR2 reconstruction from native mouse hearts. When compared to the mouse RyR2 structure obtained for the protein heterologous expressed in HEK293 cells (PDB code 7VMO) solved in the presence of Ca^2+^ ^52^, the native and heterologously expressed structures are very similar, particularly in the pore region (amino acids 4749-4961), for which the RMSD value is 0.6 Å (**Supplementary Figure 4d**).

### Nb binds to the conserved surface of the Repeat12 domain

All Nb-purified reconstructions of RyR1 and RyR2 contained additional density filling the clump-like opening of the Repeat12 domain (aa 850-1055 in rabbit RyR1, and 861-1066 in mouse RyR2) located at the corners of the square-shaped cytoplasm-exposed RyR crown (**Figures 2-5**).

The additional density was consistent with a Nb tightly docked into the crevice formed in between the helices of the repeat motifs. The polypeptide backbone was well resolved in the reconstruction of RyR1-ND which permitted to model the structure of the Nb, whereas density for the megabody’s scaffold domain cHopQ was missing, likely due to its flexibility commonly observed for the scaffold domain ^47^. Regions of the Nb interacting directly with the Repeat12 domain were particularly well resolved, whereas the density was weaker at the constant end of the nanobody. The initial model of Nb9657 generated with AF2 was manually corrected in the complementarity determining regions 1 (CDR1) and CDR3 (residues 25-33 and 96-117). Nb9657 interacts with the Repeat12 domain over an area of 1200 Å and has high shape complementarity. The interaction is primarily mediated by CDR3 that inserts deep into the opening created between the structural repeats whereas CDR1 and 2 loops provide a minor contribution (**Figure 6a**). Even though the local resolution was not sufficient to unambiguously position side chains in the interacting regions, analysis of the plausible contacts suggests that CDR3 and Repeat12 interact via multiple polar contacts involving CDR3 backbone and side chains (**Figure 6a**).

**Figure 6.**
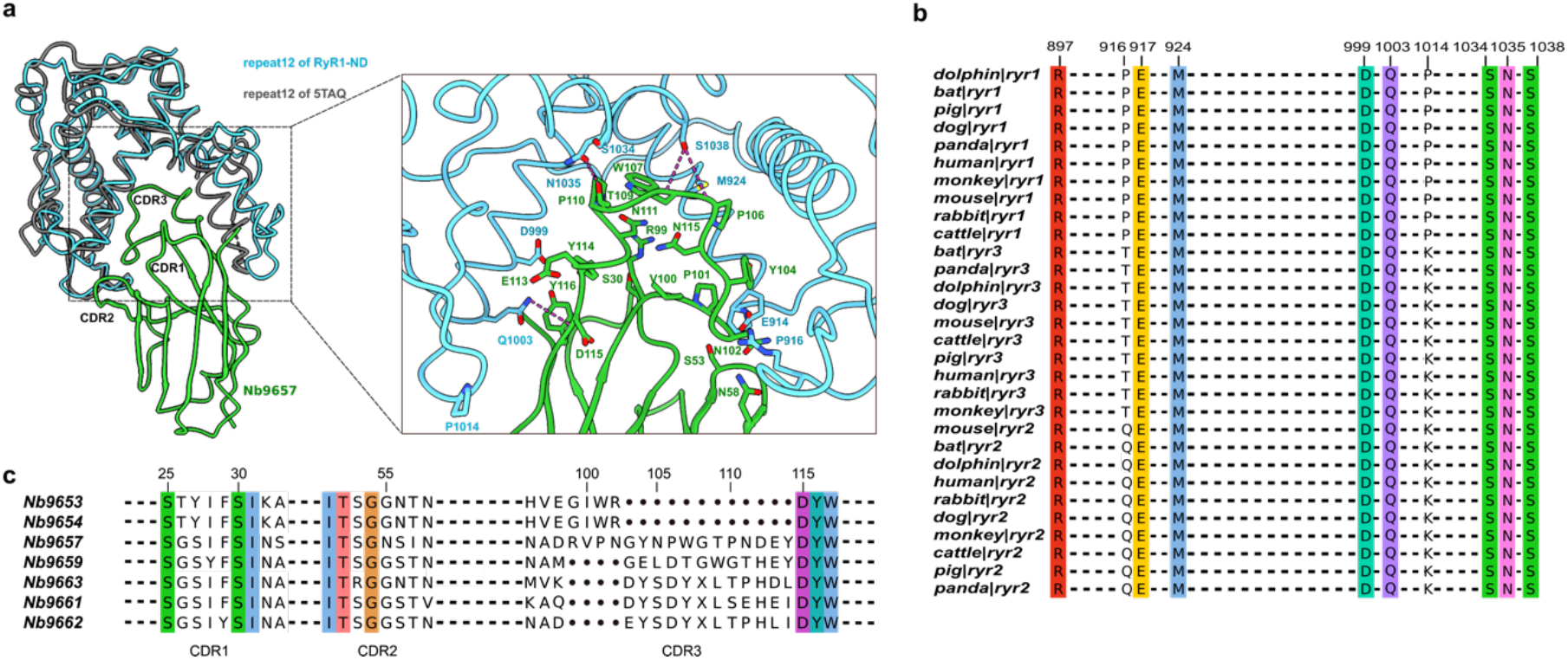
Analysis of rabbit RyR1-Nb9657 interaction. **a,** Structural alignment of the Reapeat12 domain bound to the nanobody (RyR1-ND) and free (5TAQ). **b,** Alignment of the Repeat12 domain of RyR1, RyR2 and RyR3 for 10 mammals, the residues are numbers are indicated for rabbit RyR1 sequence. Ten Repeat12 domain residues putatively interacting with the nanobody are shown, residues not involved in the interaction with the nanobody are indicated with dashes; fully conserved residues are colored. **c,** Alignment of the seven selected nanobodies. Only CDRs residues are shown, residues in between the CDRs are indicated with dashes; the dots in CDR3 parts indicate gaps in the sequence alignments.

Comparison of the structures of the Repeat12 domains with and without bound Nb9657 suggests that binding of the Nb results in the opening of the crevice through shifting of the α-helices by around 4 Å (**Figure 6a**). Interestingly, the binding site of the Nb overlaps with the binding site of the therapeutic molecule ARM210 that binds to the repeat12 domain cooperatively with ATP ^53^. Whereas the binding of Nb9657 expanded the crevice, binding of ARM210 had the opposite effect and made it tighter. Among all the selected Nbs, Nb9657 has the longest CDR3 loop (**Figure 6c**) consistent with its high affinity. At least one more nanobody, Nb9662, bound to the same binding site but in a different orientation, as can be deduced from 7.5 Å resolution map of bovine RyR2 (**Supplementary Figure 5**), suggests that the induced conformational changes might be nanobody-specific.

Furthermore, we analyzed the conservation of the Repeat12 surface residues interacting with the Nb9657 in different RyR isoforms and organisms (**Figure 6b**). Among 10 residues of rabbit RyR1 that putatively interact with Nb9657 eight are fully conserved between all three RyR isoforms suggesting that the Nb can be used to purify RyR isoforms from various mammals and will be useful for biochemical, biophysical and structural studies of mammalian RyRs, including human.

## Discussion

Here, we described the generation of high affinity nanobodies against RyR and their application to the purification of RyR1 and RyR2 isoforms in a form directly suitable for high-resolution structural analysis by single particle cryo-EM. The developed purification strategy involves trapping solubilized RyR by Nb9657 bound to magnetic beads via His-tag. This purification step is repeated twice to reach cryo-EM-grade protein purity. Even though Nb-RyR interaction is very specific, the impurities after single purification step are likely caused by poor specificity of Ni-NTA over other affinity tags based for example on streptavidin, maltose binding protein, or FLAG-peptide antibodies ^54^. The advantage of Ni-NTA beads however, is their high capacity and efficiency of protein elution, which permits maintaining high protein concentration throughout purification process thus eliminating a need for often delicate protein concentrating step.

The two affinity purification steps are followed by quick buffer exchange steps on a miniature 200 μl spin columns which are used to remove excess of Nbs or Mbs, replace buffer, and reconstitute RyR into liposomes.

Importantly, along with the purification method we report an optimized procedure for preparation of cryo-EM grids in which preferential orientation was reduced by using megabodies.

Among the core advantages of the developed method over the other described purification strategies is the possibility of purifying RyR on a small scale. RyR amounts sufficient for preparation of cryo-EM grids can be purified starting from around 5 mg of total protein contained in SR membranes. This corresponds, for example, to 15 mouse hearts, 100 mg each. Additionally, the purification procedure is fast. It takes only 4 hours starting from isolated SR membranes to cryo-EM grids. The longest step is the exchange of nanobody with megabody, which takes at least one hour. This is different from the FKBP-based approach in which natively bound FKBP is replaced by affinity tag containing FKBP, which is a slow step due to the low Koff value of FKBP ^34,35^. The purification procedure we developed is performed on a bench and does not require specialized equipment like an HPLC system or commercial prepacked columns. Furthermore, the purification is soft, it retains native FKBP bound to RyR1 (**Figures 2-5**). The buffer exchange steps included in the purification allow for a convenient way of introducing ligands and small proteins into the final purified protein and the eluent is directly suitable for cryo-EM grid plunging.

We show that essentially the same purification procedure can be easily applied to prepare RyR in various lipid mimetics, including a standard detergent-solubilized protein, MSP-based lipid nanodiscs and lipid liposomes. For all these conditions we solved RyR structures.

Interestingly, despite the presence of RyR activators in RyR1-ND sample, the channel conformation corresponded to 100% occupancy of primed state whereas under otherwise identical conditions when the same protein was purified in detergent, 16% of the particles had open channel conformation, suggesting that nanodiscs reduce the open probability of RyR. The RyR1 open probability under various lipid mimicking conditions is further investigated in a follow up study.

We selected 7 Nbs which pulled down both RyR1 and RyR2 isoforms. Of those Nb9657 and Nb9662 were the most effective (**Figure 1**). Nb9657 was used for establishing purification protocols because of its higher stability. The cryo-EM reconstructions allowed us to determine the details of the RyR1-Nb9657 interactions. The Nb tightly docks into the wedge between the helical structural repeats in the Repeat12 domain and its position is identical for RyR1 and RyR2. Repeat12-RyR interactions are mediated by an extended CRD3 loop and the interaction surface of Repeat12 is relatively well conserved between RyR isoforms and between RyRs from different organisms, suggesting that the Nb9657 will also be efficient in purifying RyR3 isoform which has not been structurally characterized yet.

We noticed that binding of Nb9657 induces conformational changes in the Repeat12 domain **(Figure 6a)** by opening the cavity between the structural repeats. This local domain modification does not have apparent effect on the structures of the investigated RyRs beyond the Repeat12 domain. Furthermore, our structural data suggest that both RyR1 and RyR2 can be opened by common activators indicating that bound to Nb9657 both RyR1 and RyR2 receptors remain functional. Nb9657 occupies the same binding site as ARM210^53^ but induces conformational changes in the opposite direction. Whilst ARM210 stabilizes RyR1 in the closed conformation, binding of the nanobody did not have obvious effect on the channel opening although electrophysiological characterization will be needed to assess possible smaller effects of the nanobody on RyR open probability.

Requirement for a small amount of starting material combined with a fast and soft purification method and conservation of the Nb9657-RyR interaction surface opens possibilities for the studies of the structures of all RyR isoforms from various mammals. In this work, we report reconstructions of RyR2 from bovine heart and mouse heart both purified from native tissues demonstrating utility of the reported purification method.

Because the described purification procedure allows for the straightforward exchange of small molecules and proteins in the solution of purified RyR, it will facilitate studies of RyR regulation by numerous interacting molecules and can also facilitate efficient small molecule screening for drug development.

## Acknowledgments

We thank Annelore Stroobants for technical support, Dr. Marcus Fislage for assistance with cryo-EM data collection and Eva Beke for assistance with the generation of the nanobody libraries. Data acquisition at the National Center for Electron Nanoscopy in Leiden (NeCEN) was co-financed by grants from the Nederlandse Organisatie voor Wetenschappelijk Onderzoek (project 175.010.2009.001) and by the European Union’s Regional Development Fund through ‘Kansen voor West’ (project 21Z.014). We acknowledge the support and the use of resources of Instruct-ERIC, part of the European Strategy Forum on Research Infrastructures (ESFRI), and the Research Foundation - Flanders (FWO) for their support to the Nanobody discovery. This work was funded by grants from FWO (grant Nos. G.0266.15N, G0H5916N, G054617N to R.G.E.), IWT (fellowship number 131261 to K.W.) and the European Research Council (Grant No. 726436 to R.G.E.)

## Data availability

The newly generated cryo-EM density maps and refined atomic models have been deposited to the PDB and EMDB databases under accession codes: PDB-8RRX and EMD-19468 for the RyR1-ND, 8RS0 and EMD-19472 for RyR1-DT-primed state, 8RRW and EMD-19467 for RyR1-DT-open state, 8RRV and EMD-19466 for RyR1-DT-closed state, 8RRU and EMD-19465 for RyR1-LP-primed state, 8RRT and EMD-19464 for RyR1-LP-open state, 8RRS and EMD-19463 for RyR2-DT. Density maps of bovine RyR2 tetramer and dimer of tetramers were deposited to EMDB with accession codes EMD-11071 – reconstruction of bovine RyR2-FKBP12.6-Nb, EMD-11072 – reconstruction of RyR2-Nb dimer.

## Contributions

C.L. optimized conditions for cryo-EM grid preparation for RyR1 and RyR2, purified protein, obtained 3D reconstructions, refined models, and wrote the original draft of the manuscript. K.W. purified RyRs, selected nanobodies, established purification procedure for RyR2, collected and processed cryo-EM data and contributed to the manuscript writing. T.U. generated the megabody. E.P. and J.S. supervised nanobody production and selection. R.G.E. supervised project, reviewed and edited the manuscript, analyzed data. K.W., J.S. and R.G.E. acquired funding.

## Methods

### Isolation of rabbit skeletal SR membranes

Sarcoplasmic reticulum membranes were isolated from rabbit fast twitch muscles using a modified protocol based on previously described method ^31^. Fresh muscle tissue (~120g) of a male New Zealand White rabbit of 8 weeks old was minced and homogenized in ice-cold RyR1 homogenization buffer (20 mM Tris-maleate pH 7.0, 100 mM NaCl, 0.3 M sucrose, 1 mM EGTA, 2 mM DTT, and a cocktail of inhibitors: 4 mM leupeptin, 1 mM benzamidine, 0.1 mM aprotinin, 1 mM pepstatin, 2 mM calpain I inhibitor and 1 mM phenylmethylsulphonyl fluoride (PMSF)) in a volume of 1.6 l using a Waring blender (3 X 30 s). The homogenized suspension was centrifuged for 30 min at 7,000 rpm in Beckman JA-10 rotor at 4°C. Crystalline KCl was added to the supernatant filtered through cheesecloth to a final concentration of 0.5 M and stirred between 10 and 90 min at 4°C. The extract was centrifuged at 40,000 rpm in Beckman 45Ti rotor for 30 min at 4°C. The resulting pellet was homogenized in resuspension buffer (20 mM MOPS, pH 7.4, 1 mM EGTA, 2 mM DTT, 0.6 M KCl, 0.3 M Sucrose and cocktail of inhibitors) in a volume of 80-100 ml with a glass homogenizer and centrifuged at 40,000 rpm in Beckman 45Ti rotor for 30 min at 4°C. The pellets were homogenized in resuspension buffer at total protein concentration of around 10 mg/ml, frozen in liquid nitrogen and stored at -80°C. Protein concentration was determined by Pierce 660 nm Protein assay (Thermo Fisher Scientific).

### Isolation of mouse cardiac SR membranes

Mouse cardiac SR membranes were prepared following protocol described elsewhere ^31^, with following modifications. Fresh mouse hearts were collected after the animals were sacrificed and embedded in ice cold RyR2 homogenization buffer (25 mM Hepes pH7.4, 150 mM NaCl, 0.5 mM EGTA, 10% sucrose, 2 mM TECP and a cocktail of inhibitors). Two hundred mouse hearts were homogenized in 400 ml of homogenization buffer in a blender and centrifuged in Beckman JA-20 rotor in 7,000 rpm for 30 min. The following steps were the same as for the isolation of rabbit skeletal SR membranes with modified buffer composition. The RyR2 resuspension buffer was similar to RyR2 homogenization buffer except for the NaCl concentration of 1 M. After the last ultracentrifugation, the pellets were resuspension in 30 ml of resuspension buffer at a concentration of around 8 mg/ml, frozen in liquid nitrogen and stored at -80°C. Protein concentration was determined by Pierce 660 nm Protein assay (Thermo Scientific).

### Isolation of bovine cardiac SR membranes

Bovine cardiac SR membranes were prepared using protocols described above, with the following modifications. Fresh bovine heart was collected from a slaughterhouse (approximately 20 min postmortem) and transported within 30-60 min to the laboratory in ice-cold cardioplegic buffer (25 mM Tris/Hepes pH 7.4, 102 mM NaCl, 21.4 mM KCl, 16 mM MgCl2, 2.4 mM CaCl2). Ventricles were separated from fat, connective tissue, and blood vessels, and cut in small pieces. Ventricles from one heart (∼1 kg) were homogenized in 3 l of buffer B (25 mM Tris/Hepes pH7.4, 0.3 M sucrose and a cocktail of inhibitors). Pellets from the first round of ultracentrifugation were homogenized in buffer C (buffer B supplemented with 400 mM KCl, 0.5 mM MgCl2, 0.5 mM CaCl2, and 0.5 mM EGTA) to a final volume of 420 ml with a glass homogenizer. The resuspended membranes were spun down at 130,000 g for 30 min at 4°C. Next, pellets were homogenized in buffer C at a concentration of 15-20 mg/ml, frozen in liquid nitrogen and stored at -80°C. Protein concentration was determined by Pierce 660 nm Protein assay (Thermo Fisher Scientific).

### Immunization and selection of nanobodies

Rabbit RyR1 was purified following a previously described procedure ^55^ and reconstituted into amphipol A8-35 by mixing at 1:10 (w/w) with RyR1 followed by overnight detergent removal with BioBeads (Bio-Rad). For the Nb selection, bovine RyR2 was crudely purified from cow heart in detergent with lipids (0.6% CHAPS, 0.3% soybean polar lipids) on sucrose gradient.

The immunization, nanobody selection and screening were performed as described elsewhere ^56^. One llama (*Lama glama*) was immunized six times with a total amount of 0.87 mg of rabbit RyR1 reconstituted in amphipols (RyR1-amp). A phage immune library was generated. Phage display selections were performed on (1) RyR1-amp in 10% sucrose 200 µM EGTA 2 mM DTT, (2) on RyR1 in 20 mM MOPS 0.7 M NaCl ∼20% sucrose 200 µM EGTA 2 mM DTT 200 µM CaCl2 2 mg/ml soybean lipids 0.8% CHAPS and (3) on RyR2 in 20 mM MOPS 0.7M NaCl ∼20% sucrose 200 µM EGTA 2 mM DTT 200 µM CaCl2 2 mg/ml soybean lipids 0.8% CHAPS. The sequence of 276 single colonies (92 colonies picked from each condition) was analyzed and 106 Nbs sequences were grouped according to the CDR3 sequence into 13 families whereas the remaining 170 Nbs were single representatives. From the 13 families, 38 Nbs were screened in ELISA using periplasmic extracts for binding to rabbit RyR1. To avoid selecting Nbs against SERCA1a (sarcoplasmic/endoplasmic reticulum calcium ATPase) which was a significant contaminant of purified RyR1 used for immunization, sample containing enriched SERCA1a fractions from size-exclusion chromatography on Superdex 200 obtained during RyR1 purification was used as a negative control. Nbs producing an ELISA signal twice higher than the signal of the negative control wells were retained (**Supplementary Figure 1a**). This identified 25 binders for skeletal RyR1 out of which 17 nanobodies belonging to 10 different families were screened using ELISA for binding to crudely purified solubilized in CHAPS bovine RyR2 which gave significant signal for 7 Nbs (**Supplementary Figure 1b**) which were retained for further purifications.

### Preparation of periplasmic extract containing Nbs

The nanobody repertoire was cloned into phage-display vector pMESy4 which allows for expression of Nbs with a 6xHis-tag. Overexpression of Nbs was done overnight after induction by 1 mM Isopropyl β-d-1-thiogalactopyranoside (IPTG) of *Escherichia coli* WK6 strain after the cell density reached OD of 0.8 at 600 nm. Cells are collected by a 15 min centrifugation at 5,000 rpm in Beckman JA-10 rotor. Cells were lysed by adding 4 mL of lysis buffer [50 mM Tris pH 7.5, 150 mM NaCl, 1 mM EDTA, 0.1 mg/ml lysozyme, 20% sucrose, 50 µg/ml DNase, and 1 mM phenylmethylsulphonyl fluoride (PMSF)] per 1 g of pellet followed by 30 min incubation at 4°C. Periplasmic extract was obtained by 15 min centrifugation at 10,000 rpm in Beckman JA-20 rotor. MgCl2 (5 mM) was added to the extract, the extract was filtered, supplemented with 20% sucrose, and stored at -80°C in aliquots.

### Megabody purification

The Mb9657 was constructed by grafting Nb9657 onto scaffold protein cHopQ with His6 tag on C-terminus following the protocol described elsewhere ^47^. The megabody was expressed overnight in *Escherichia coli* WK6 strain after induction with 1 mM IPTG at OD between 2 and 3 at 600 nm. Cells were collected by a 15 min centrifugation at 5,000 rpm in Beckman JA-10 rotor. Periplasmic extract was obtained by homogenizing an equivalent of 1 l cell culture pellet in 25 mL of lysis buffer (50 mM Tris pH 8, 150 mM NaCl, 1 mM EDTA, 0.3 mg/ml lysozyme, 20% sucrose, 1 µg/ml leupeptin and 0.1 mg/ml AEBSF) followed by 30 min incubation at 4°C. Next, NaCl concentration was increased to 500 mM NaCl, 5 mM MgCl2 and 50 µg/ml DNase were added, cells were centrifuged at 10,000 rpm in Beckman JA-20 rotor for 1 h and resulting supernatant was passed through 0.45 µm filter.

The periplasmic extract was then loaded on the 1 ml HisTrap HP column pre-equilibrated in 10 column volumes (CV) of wash buffer (500 mM NaCl, 100 mM Tris-HCl, 5 mM imidazole) at a flowrate of 5 ml/min, then washed by 10 CV of wash buffer and eluted in 3 CV of elution buffer (similar as wash buffer except for the imidazole concentration of 500 mM). The sample was concentrated on a spin column (10 kDa, Meck) to 1 ml and loaded on a Superdex 200 Increase 10/300 GL column equilibrated with gel filtration buffer (10 mM Tris-HCL, 140 mM NaCl, PH 7.4) and eluted with 1.2 CV. The peak fractions were collected and concentrated on a spin column (10 kDa, Meck) to 14.8 mg/ml.

### Pull-down assays for RyR1

Rabbit skeleton SR membranes 0.5 ml were solubilized in 375 µl of solubilization buffer (20 mM MOPS pH7.4, 1 M NaCl, 10% sucrose, 2 mM TECP, 4% CHAPS 0.8% POPC and a cocktail of inhibitors) over 30 min at 4°C. Detergent solubilized membranes were spun at 40,000 rpm in Beckman 50.4 Ti rotor for 30 min. 5 µl of HisPur™ Ni-NTA Magnetic Beads (MB, Thermo Fisher Scientific) were loaded with Nb by incubation with 500 µl of periplasmic extract for one hour at 4°C on a rotator. Next, MB were washed in equilibration buffer (20 mM MOPS pH7.4, 1M NaCl, 10% sucrose, 2 mM TECP, 0.8% CHAPS 0.2% POPC and a cocktail of inhibitors). The Nb-loaded MB were incubated with different dilutions of the detergent solubilized membranes: non-diluted, x5 and x25 in equilibration buffer. The total amount of protein incubated with the MB was equal for every condition. The samples were incubated for one hour at 4°C while steering. After washing with equilibration buffer, MB were loaded on SDS-PAGE (4–15% Mini-PROTEAN® TGX™ Precast Protein Gels, Bio-rad). Gels were stained with Coomassie-blue.

### Pull-down assays for RyR2

Cardiac SR membranes were solubilized by 1:1 dilution in solubilization buffer (25 mM PIPES pH 7.4, 1 M NaCl, 150 µM CaCl2, 100 µM EGTA, 0.6% CHAPS, 0.3% soybean polar lipids, and protease inhibitors) over 2 hours at 4°C. Detergent solubilized membranes were spun at 103 000 g for 30 min. The remaining procedures were similar to RyR1 pool down assays except for the equilibration buffer composition (25 mM PIPES pH 7.4, 0.3 M NaCl, 0.6% CHAPS, 0.3% soybean polar lipids).

### Purification of membrane scaffold protein MSP1E3D1

Membrane scaffold protein MSP1E3D1 (6xHis-tagged) was purified as described elsewhere ^57^ with some modifications. MSP1E3D1 was overexpressed in *E. coli* BL21 (DE3) with 1 mM IPTG. Cells pellets were collected and dissolved in 20 mM NaPi pH 7.4 and disrupted with a high-pressure homogenizer (Constant Systems Ltd). The cell debris were removed by centrifugation, the MSP1E3D1 was purified on a 5 ml HisTrap™ High Performance column GE Healthcare). MSP was eluted in buffer (40 mM Tris pH 7.5, 0.3 M NaCl, 1% Na Cholate with 400 mM Imidazole) and dialyzed overnight against a dialysis buffer (40 mM Tris pH 7.5, 100 mM NaCl). CHAPS was added to the dialyzed sample to concentration of 1% after which the MSP was concentrated to 20-40 mg/ml in centrifugal filter (Merck, MWCO 30 kDa), flash frozen and stored at -80°C.

### Purification and reconstitution of RyR1 into lipid nanodiscs

Skeletal SR membranes (0.5 ml, 10 mg/ml) were solubilized in 375 µl of solubilization buffer (20 mM MOPS pH 7.4, 1 M NaCl, 10% sucrose, 2 mM TECP, 4% CHAPS, 0.8% POPC, and a cocktail of protease inhibitors). Every step of the purification was performed at 4°C or on ice. The sample was incubated for at least 30 min and ultracentrifuged for 30 min at 40,000 rpm in Beckman 50.4 Ti rotor. Magnetic beads were washed 3 times with 270 µl SBL buffer (similar to solubilization buffer except for 0.7 M NaCl, 0.8% CHAPS and 0.2% POPC, and 200 µM EGTA). MB were loaded with Nb9657 by incubating 135 µl Nb9657 periplasmic extract with 135 µl equilibrated magnetic beads in 1.5 ml eppendof tubes for 15 min). Nb-loaded MB (135µl) were incubated with 800 µl (5mg total protein) of detergent-solubilized membranes for 30 min and washed with SBL buffer. RyR was eluted by incubating MBs in 65 µl of elution buffer 1 (SBL buffer with 500 mM imidazole) for 5 min. The eluate was diluted three-fold in SBL buffer to lower the imidazole concentration. MSP1E3D1 was added at a 1:130 MSP:phospholipid molar ratio and incubated for 15 min. RyR1 was reconstituted into lipid nanodiscs by diluting the detergent with dilution buffer (SBL buffer without CHAPS and POPC) three times over 30 min using a syringe pump. Next, 0.2% fOM was added and incubated for 20 min. For the second purification, another batch of 135 µl Nb9657-loaded MB was incubated with the reconstituted sample for 30 min, then washed with fOM buffer (dilution buffer with 0.2% fOM). Reconstituted RyR1 was eluted by incubating the magnetic beads for 5 min in 50 µl of elution buffer 2 (fOM buffer with 2 mM ATP, 5 mM caffeine, 250 µM CaCl2 (corresponds to 50 µM free Ca^2+^) and 500 mM Imidazole).

For the samples without megabody, the first desalting was done on a homemade desalting column containing 400 µl of Superdex200 prep grad resin (Cytiva) loaded in a 1.5 ml spin column for protein isolation (Carl Roth). The column was pre-equilibrated with desalting buffer 1 (similar as elution buffer 2, except the fOM concentration of 0.06% and NaCl concentration of 0.2 M). 50 µl of RyR collected after second elution was applied on the column and centrifuged at 1000 x g for 1min. The desalted sample was used for plunging.

For sample with Mb9657, desalting was repeated twice. The first desalting was done as described above, except the fOM concentration in desalting buffer changed to 0.2%. Eluted protein was incubated with 60 µg of Mb9657 for at least one hour followed by a second round of desalting in the column pre-equilibrate in desalting buffer 1.

### Purification of RyR1 solubilized in detergent

The purification of RyR1 in detergent micelles was identical to the previous procedure up to and including the first elution step. Then the 65 μl of eluate 1 was diluted with 600 µl of SBL buffer and incubated 30 min with 135 µl of Nb9657 loaded MB. MB were washed and RyR1 was eluted directly after the incubation following procedure described above with modified buffers. For preparation of RyR1 without activators, the elution buffer 2 was SBL buffer with 500 mM imidazole. The composition of the desalting buffer 1 was similar to the elution buffer 2 but imidazole was omitted. The desalting buffer 2 composition was similar to the desalting buffer 1 with fOM replaced by 0.2% CHASP and 0.001% POPC. Preparation of RyR1 with activators followed the same procedure with 2 mM ATP, 5 mM caffeine and 250 µM CaCl2 (50 µM free Ca^2+^) added to the elution buffer 2 and the desalting buffers.

### Purification and reconstitution of RyR1 into liposomes

The purification procedure followed the same steps as the RyR1 purification in detergent till and including the second elution. After second elution, one desalting step was done with buffer similar to desalting buffer 1 with fOM replaced with 0.008% CHAPS and 0.01% POPC.

### Purification of mouse RyR2 solubilized in detergent

RyR2 solubilized in CAHPS was purified like RyR1 solubilized in CHAPS with the following modifications. During solubilization, heart SR membranes (1.6 ml, 8 mg/ml) were solubilized in 1.6 ml of RyR2 sol buffer (50 mM HEPES, 1 M NaCl, 5 mM TECP, 0.6% CHAPS, 0.3% POPC and cocktail of protease inhibitors). Other steps were the same as for as RyR1 except for the SBL buffer composition which was the same as the RyR2 sol buffer. The first desalting buffer was RyR2 sol buffer with 2 mM ATP, 5 mM caffeine, 50 µM Ca^2+^. The second desalting buffer was similar to the first but contained 0.2% CHPAS and 0.001% POPC.

### Purification of bovine RyR2 reconstituted into lipid nanodiscs

Bovine RyR2 purification was similar to the purification of RyR1-ND, with some modifications. Cardiac SR membranes (~16mg/ml) were solubilized by 1:1 dilution with solubilization buffer (25 mM Tris/50 mM HEPES pH7.4, 1 M NaCl, 0.6% CHAPS, 5 mM TCEP, and protease inhibitors). MB were equilibrated washed with SBL buffer (25 mM Tris/50 mM HEPES pH7.4, 1 M NaCl, 0.6% CHAPS, 0.3% soybean polar lipids, 20% sucrose, 5 mM TCEP and protease inhibitors). 100 µl of MB was used for the purification of 1.5 ml detergent-solubilized membranes. The Nb-loaded MB were incubated with the solubilizate 30 min and washed with SBL buffer. Elution was done in SBL buffer with 500 mM imidazole. The reconstitution of bovine RyR2 into nanodisc and latter incubation with fOM was as for RyR1-ND. Purified RyR2 was dialyzed to fOM buffer in Slide-A-Lyzer™ MINI Dialysis unit 20K MWCO 0.1 mL (Thermo Scientific) overnight. After dialysis, the protein is used immediately for for cryo-EM sample preparation.

### Preparation of graphene oxide grids

Home-made graphene oxide grids were used for plunging. A stock solution was prepared by diluting 2 mg/ml graphene oxide solution (Sigma with Milli Q water to 0.6 mg/ml). Every time before using, the stock solution was centrifuged for 3 min at 3000 g and the supernatant was diluted 5 times with milli Q water to 0.12 mg/ml. Quantifoil R2/1 Cu300 holey carbon grids were glow discharged in the ELMO glow discharge system (Corduan Technologies) from carbon side for 1 min at 5 mA and 0.28 mbar. A volume of 3µl of 0.12 mg/ml graphene oxide was applied to the carbon side and incubate for at least 1 min. The grids were blotted with Whatman filter paper No. 1 and washed with 10 µl distilled water and blotted immediately with Whatman No. 1 filter paper. The grids were dried for at least 30 min at room temperature.

### Preparation of cryo-EM grids

The cryo-EM samples were prepared using a CP3 cryoplunge (Gatan). The protein solution (4 µl) was loaded on the carbon side, 1 µl of desalting buffer without fOM or lipids (depending on the sample) was loaded on the back side and blotted from both sides for 1.2 s with Whatman glass microfiber filters paper GF/A at 91% relative humidity. The grids were plunge-frozen in liquid ethane at −175°C and stored in liquid nitrogen.

### Cryo-EM data collection

Single particle cryo-EM data for bovine RyR2-FKBP12.6-Nb were collected at NeCEN (University of Leiden, Netherlands) on a Titan Krios equipped with a Falcon III using automated data collection software, EPU (FEI). A total of 4,088 usable micrographs were collected in linear mode at a nominal magnification of 59,000 and corresponding pixel size of 1.4 Å with 29 frames and a total dose of 47 e-/Å^2^. Images were recorded with defocus between -2 and -4 µm.

All the remaining cryo-EM images were collected on a JEOL CryoARM 300 microscope equipped with an in-column Ω energy filter, a cold field emission gun (cFEG) ^58^ operating at 300 kV. K3 detector (Gatan) was used to record images. Movies consisting of 60 frames at nominal magnification of 60,000 were collected automatically using SerialEM 3.0.8 ^59^. The energy filter slit was set to 20 eV width. The exposure time was 2.796 s with average dose per frame of 1 e^-^/Å^−2^ and defocus values in the range of -1.5 to -2.5 µm. The calibrated pixel sizes were between 0.73 Å to 0.76 Å. Five images per single stage position were collected using a cross pattern with three holes along each axis ^60^.

### EM image processing

For all datasets, motion correction was done using MotionCor2 ^61^. The Contrast Transfer Function (CTF) parameters were estimated using CTFFIND-4.1 ^62^. The images were imported to cryoSPARC v 4.2.1 or v 3.3.2^63^ for further processing. Particle picking was done by template picker with 2D templates created with particles picked manually or created from already processed datasets. After particle picking, all particles were extracted in 4-times binned box with 672 pixels. Then several rounds of 2D classification were performed to select particles for further processing see (**Supplementary Figures 7-11**).

For RyR1-ND dataset, 540,520 particles were selected after 2D classification and used to generate an initial model with C1 symmetry. 537,558 particles were re-extracted in box size 672 pixels binned twice refined to a consensus 3D map with C4 symmetry. Then particles with refined orientation parameters were import to Relion v3.1.3 ^64^ for 3D classification into 8 classes with a mask around the transmembrane part (**Supplementary Figure 7**) to separate open and closed states. The channel diameter was the same for all the classes and corresponded to the closed conformation. To improve the resolution, classes 5,6 and 8 with the highest resolution accounting for 175,535 particles were joined, re-extract in a box with 450 pixels Fourier cropped to 1.49 Å and a consensus 3D map was refined to 3.1 Å. Then Bayesian polishing was performed in Relion v 4.0.0. The polished particles were refined in CryoSPRC with C4 symmetry and per-particle defocus, per-group CTF parameters, fit beam tilt, beam trefoil, spherical aberration, beam tetrafoil and beam anisotropic magnification. This resulted in a consensus map resolved to 3.1 Å. To improve the density in the flexible domains, four local refinements with local masks (**Supplementary Figure 7**): mask 1 covered TaF, TM and CTD, mask 2 covered JSola and CSol, mask 3 covered FKBP and TTD, mask 4 covered BSol were performed. The local refinement with mask 1 was performed on 175,535 particles with C4 symmetry. With masks 2, 3 and 4 local refinement was performed with C1 symmetry after C4 symmetry expansion. The Local maps with resolution of 2.9 Å for map 1, 2 and 3, and 3.5 Å for map 4 were produced. The resulting maps were combined in Chimera v1.15 to generate the composite map, the pixel size calibration and model building was done on the composite map.

Other datasets were processed the same way.

For RyR1-DT without activators, 171,023 particles were selected after 2D classification and used to generate ab initio model with C1 symmetry (**Supplementary Figures 9**). The particles were re-extracted in the box binned twice with 1.46 Å pixel size and refined to a consensus 3D map with C4 symmetry. All particles were imported to Relion v3.1.3 for polishing and imported back to CryoSPARC v 4.2.1. A consensus map of 3.2 Å was generated, local refinement was performed in the same way for the RyR-ND data set. Local maps with resolution of 3.0 Å for map 1 and map 2, 3.0 Å for map 3, 3.6 Å for map 4 were obtained.

For RyR1-DT with activators (**Supplementary Figures 8**), 177,251 particles were re-extracted after selection by 2D classification. The particles were polished in Relion v3.1.3 and 3D classified into 6 class with mask covering TM domain to separate open and primed states. Classes 2,4,5 corresponded to primed conformation and accounted for 145,830 particles, class 6 with 29,246 particles corresponded to open conformation. The open and primed state particles were imported back to cryoSPARC v4.2.1 and refined by NU-Refinement procedure separately. The consensus maps of open state and primed state were reconstructed to resolution of 4.2 Å and 3.3 Å, respectively. After local masked refinements, the resolution of open state local maps was 3.8, 3.9, 4.0 and 6.3 Å for maps 1-4, respectively. The resolution of primed state local maps was 3.1 Å for maps 1-3 and 3.6 Å for map 4.

For RyR1-LP, three data sets were merged and processed together. After several round of 2D classification, only particles inside liposome were selected for further processing (**Supplementary Figures 10**). In total 51,740 particles were re-extracted with pixel size 1.46 Å. The particles were polished in Relion v3.1.3 and 3D classification into 6 classes with mask around the TM domain part to separate open and closed states. Classes 1 and 5 accounting for 26,815 particles corresponded to open state. Class 6 with 16,530 particles corresponded to primed state. The particles were imported back to CryoSPARC v4.2.1 for NU-Refinement which produced the consensus map of open state at resolution of 4.6 Å and primed state at 4.7 Å. Local masked refinements of open state resulted in maps resolved to 4.0 Å (map 1), 4.5 Å (map 2 and 3) and 6.3 Å (map 4). The resolution of local maps for closed states were 4.7 Å (maps 1, 2), 5.5 Å (map 3) and 6.6 Å (map 4).

For RyR2-DT, 134,904 particles were re-extracted after 2D classification and used to generate an initial model with C1 symmetry (**Supplementary Figure 11**). The particles were re-extracted in a box of 672 pixels binned twice and refined to a consensus 3D map with C4 symmetry to resolution of 3.7 Å. The downstream processing was similar to the dataset processing described above. After 3D classification, all the well-resolved classes corresponded to open state. 76,852 particles from the best resolved classes 1 and 3 were pooled and refined in CryoSPARC v4.0.0 in C4 symmetry using NU-Refinement. This produced a consensus map resolved to 3.4 Å. The local maps were resolved to 3.2 Å (map 1 and 2), 3.4 Å (map 3) and 4.1 Å (map 4).

For the bovine RyR2 418,936 particles were automatically picked using CrYOLO ^65^. Following two cycles of 2D classification in CryoSPARC v2.12.4 and manual selection 41,112 particles were retained for homogenous refinement with C4 symmetry in CryoSPARC which resulted in a reconstruction to a resolution of 7.5 Å (**Supplementary Figure 12, Supplementary Table 2**). To reconstruct the dimer of RyR2-ND, 149,707 dimers were automatically picked using CrYOLO. The initial model was created using RVIPER in SPHIRE from 2D classes obtained using ISAC 2D classification in SPHIRE ^66^. After initial 3D classification with C1 symmetry in RELION3.0 a homogeneous class containing 39,649 particles was selected. Further classification separated data into single tetramers, distorted dimers, and symmetric dimers. Final reconstruction was obtained with selected 9,977 particles applying D4 symmetry at resolution of 13 Å (**Supplementary Figure 13**).

### Model building and refinement

A model of rabbit RyR1 (PDB: 7TZC) was rigid body fitted to each map in ChimeraX v1.7.1 ^67^ then manually rebuilt in Coot 0.98 ^68^ followed by real-space refinement in Phenix 1.20.1 ^69^ with default parameters. For RyR2, PDB with accession code 7VMO was used as initial model. Structural fragments corresponding to residues 2916-3230 and 3231-3579 were predicted in AlphaFold2 ^70^. The models were rigid body fitted into the cryo-EM maps using ChimeraX v1.7.1, manually rebuilt in Coot and refined in Phenix like RyR1 models. All models were validated with MolProbity ^71^.

