## Supplementary Information for "Rapid small-scale nanobody-assisted purification of ryanodine receptors for cryo-EM"

**Supplementary Table1** Statistics of cryo-EM data and models.

| **Sample** | **RyR1-DT** | | | **RyR1-ND** | **RyR1-LP** | | **RyR2-DT** |
|---|---|---|---|---|---|---|---|
| State | Closed | open | primed | primed | Open | Primed | Open |
| Ligands | / | | | 50µM free $Ca^{2+}$, 2mM ATP, 5mM Caffeine | | | |
| PDB ID | 8RRV | 8RRW | 8RS0 | 8RRX | 8RRT | 8RRU | 8RRS |
| EMBD ID | 19466 | 19467 | 19472 | 19468 | 19464 | 19465 | 19463 |
| | | | | **Data collection** | | | |
| Microscope | | | | JOEL CRYOARM300 | | | |
| Detector | | | | Gatan K3 | | | |
| Voltage(kV) | | | | 300 | | | |
| Magnification | | | | 60,000 | | | |
| Exposure time (S) | | | | 2.796 | | | |
| Electro dose (e -/Å $^2$) | | | | 60 | | | |
| Number of frames | | | | 60 | | | |
| Defocus range (µm) | | | | 1.5-2.5 | | | |
| Pixel size (Å) | | | | 0.76 | | | |
| Symmetry | | | | C4 | | | |
| Collected images | 11,584 | 8,069 | | 14,040 | 24,465 | | 8,276 |
| Used images | 5,891 | 5,191 | | 12,491 | 14,419 | | 4,918 |
| Final particles (N) | 171,023 | 29,246 | 145,830 | 166,010 | 26,815 | 16,530 | 76,852 |
| Global map resolution (Å) | 3.2 | 4.2 | 3.3 | 3.1 | 4.6 | 4.7 | 3.4 |
| Local maps resolution range (Å) | 3-3.6 | 3.8-6.3 | 3 -3.6 | 2.9-3.5 | 4-6.3 | 4.1-6.6 | 3.2-4.1 |
| **Model refinement** | | | | | | | |
| Refinement package | | | | Phenix v 1.20.1 | | | |
| Initial model used | | | | 7TZC | | | 7VMO |
| Model resolution [Å$^2$], FSC=0.5 | 3.3 | 7.6 | 3.4 | 3.1 | 7.4 | 8.3 | 3.5 |
| **Model composition** | | | | | | | |
| Non-hydrogen protein atoms | 142,994 | 143,720 | 143,554 | 144,120 | 143,716 | 143,338 | 136,400 |
| Protein residues | 180,88 | 182,04 | 181,62 | 182,04 | 182,04 | 182,04 | 170,72 |
| Ligands | 4 | 16 | 16 | 20 | 16 | 16 | 16 |
| **B-factors mean [Å$^2$]** | | | | | | | |
| Protein | 112.26 | 275.77 | 108.21 | 101.71 | 302.55 | 342.81 | 127.59 |
| Ligand | 99.00 | 249.25 | 95.30 | 97.78 | 344.12 | 285.80 | 63.39 |
| **R.M.S deviations** | | | | | | | |
| Bond lengths (Å) | 0.003 | 0.003 | 0.002 | 0.002 | 0.003 | 0.003 | 0.003 |
| Bond angles (°) | 0.590 | 0.631 | 0.631 | 0.558 | 0.643 | 0.663 | 0.617 |
| **Validation** | | | | | | | |
| Molprobity score | 1.66 | 1.80 | 1.69 | 1.47 | 1.81 | 1.91 | 1.77 |
| Clashscore | 11.48 | 15.26 | 12.04 | 8.81 | 15.82 | 15.88 | 13.32 |
| Poor rotamers (%) | 0.08 | 0.00 | 0.10 | 0.18 | 0.00 | 0.00 | 0.24 |
| **Ramachandran plot** | | | | | | | |
| Favored (%) | 97.61 | 97.43 | 97.54 | 98.07 | 97.47 | 96.74 | 97.30 |
| Allowed (%) | 2.37 | 2.53 | 2.46 | 1.93 | 2.46 | 3.26 | 2.68 |
| Disallowed (%) | 0.02 | 0.04 | 0.00 | 0.00 | 0.07 | 0.00 | 0.02 |

**Supplementary Table** 2 Statistics of cryo-EM data for bovine RyR2 dataset.

| **Data collection** | **RyR2 tetramers** | **Dimers of RyR2 tetramers** |
|---|---|---|
| EMBD ID | EMD-11072 | EMD-11072 |
| Electron microscope | Titan Krios | |
| Electron detector | Falcon III, linear mode | |
| Voltage (kV) | 300 | |
| Defocus range (μm) | 2 - 4 | |
| Pixel size (Å) | 1.4 | |
| Electron dose (e- $Å^{-2}$) | 47 | |
| Images | 4,088 | |
| 3D reconstruction | | |
| Final particles | 41,112 | 9,977 |
| Applied symmetry | C4 | D4 |
| Global map resolution (Å) | 7.5 | 13 |

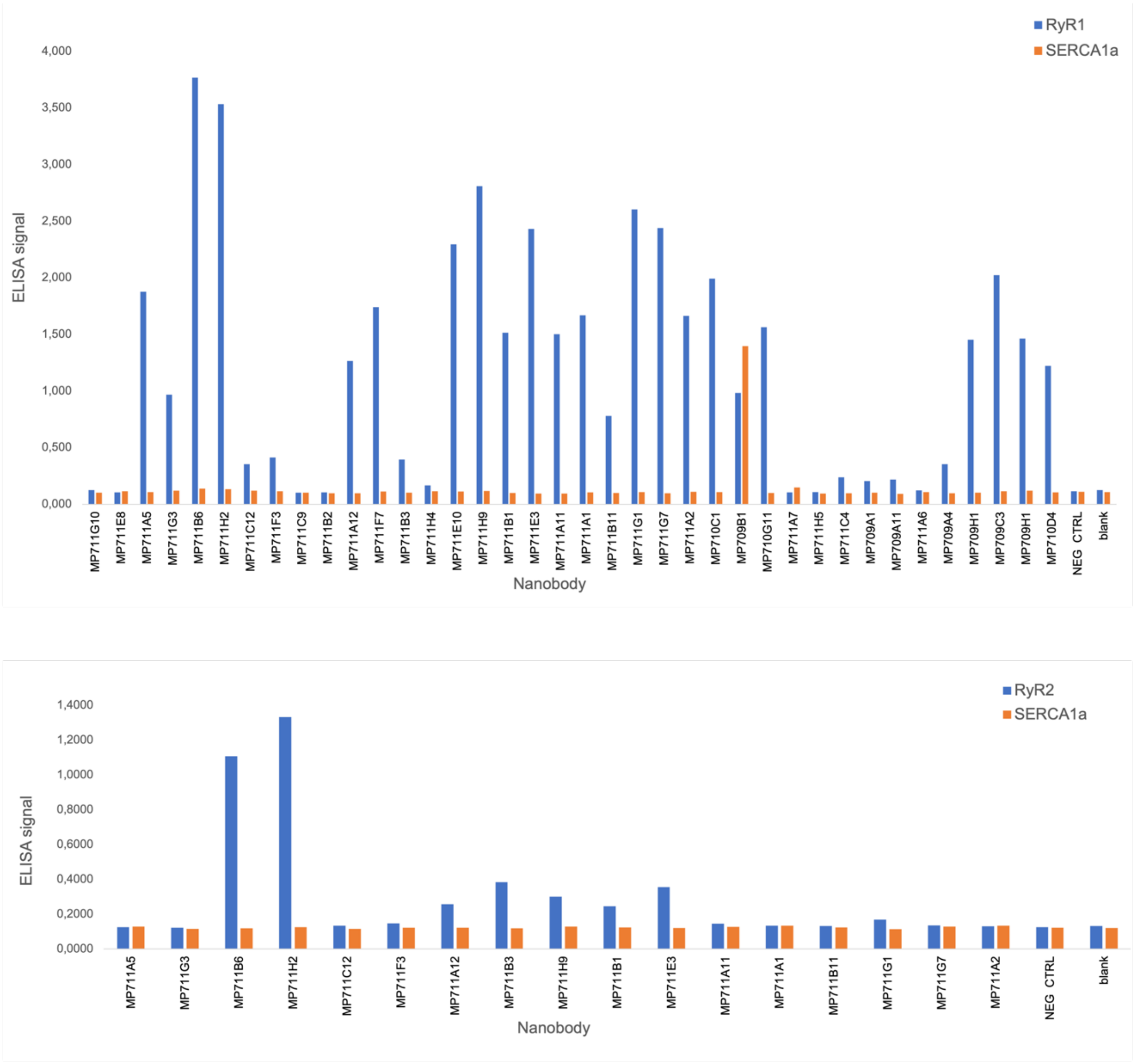


**Supplementary Figure 1 | Nanobody screening by ELISA. a,** ELISA assays on 38 selected Nbs against RyR1. For negative control solution enriched with SERCA1a was used (orange), n=1. **b,** ELISA assays on 17 Nbs against RyR2. The Nbs were picked from the pool of Nbs that reacted against RyR1. Blank signal was measured without nanobody, n=1. Seven selected Nbs are further referred to with their plasmid ID rather than clone ID with the following name mapping: MP711B6-Nb9653, MP711H2-Nb9654, MP711A12-Nb9657, MP711B3-Nb9659, MP711H9-9661, MP711B1-9662, MP711E3-Nb9663.

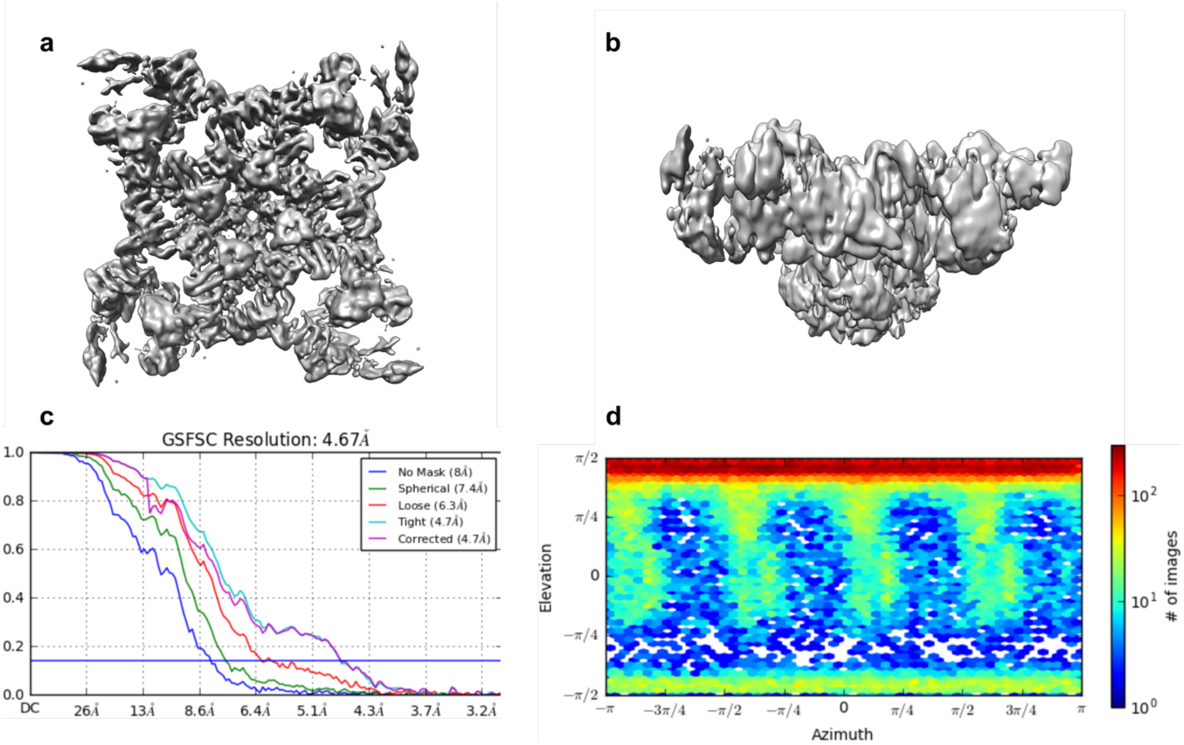


**Supplementary Figure 2 |Cryo-EM reconstruction of RyR1-ND with bound nanobodies shows strong preferred orientation.** **a**, **b** Top and side view of the cryo-EM map. **c,** FSC curves. **d,** The particle orientation distribution plot.

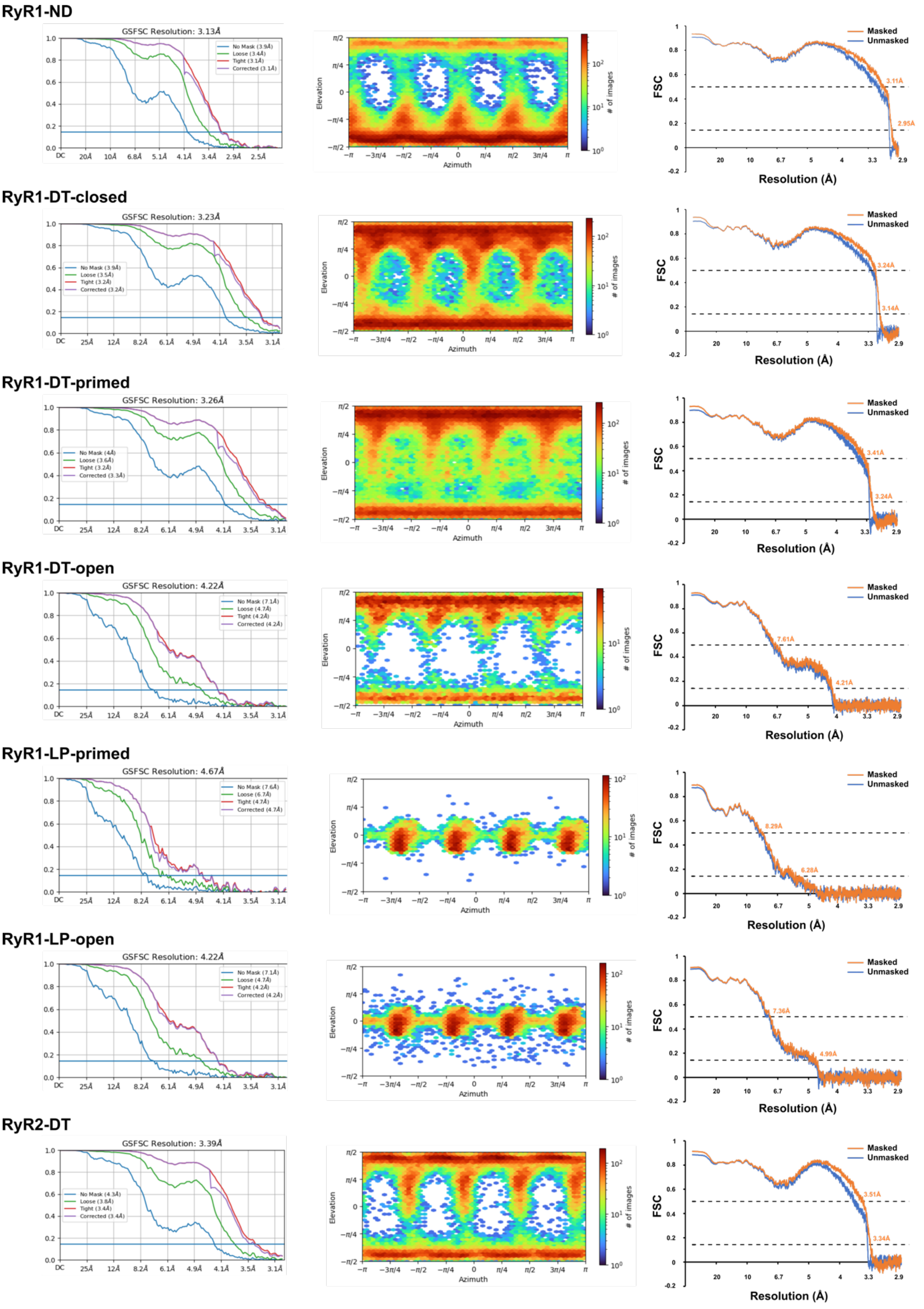


**Supplementary Figure 3 | FSC curves and distribution of orientation for rabbit and mouse RyR datasets.** FSC curves (left) and particle distributions (middle) are shown for the global non-uniform refinement performed in CryoSPARC. Model-map FSC (right) between the composite maps and refined models calculated in PHENIX.

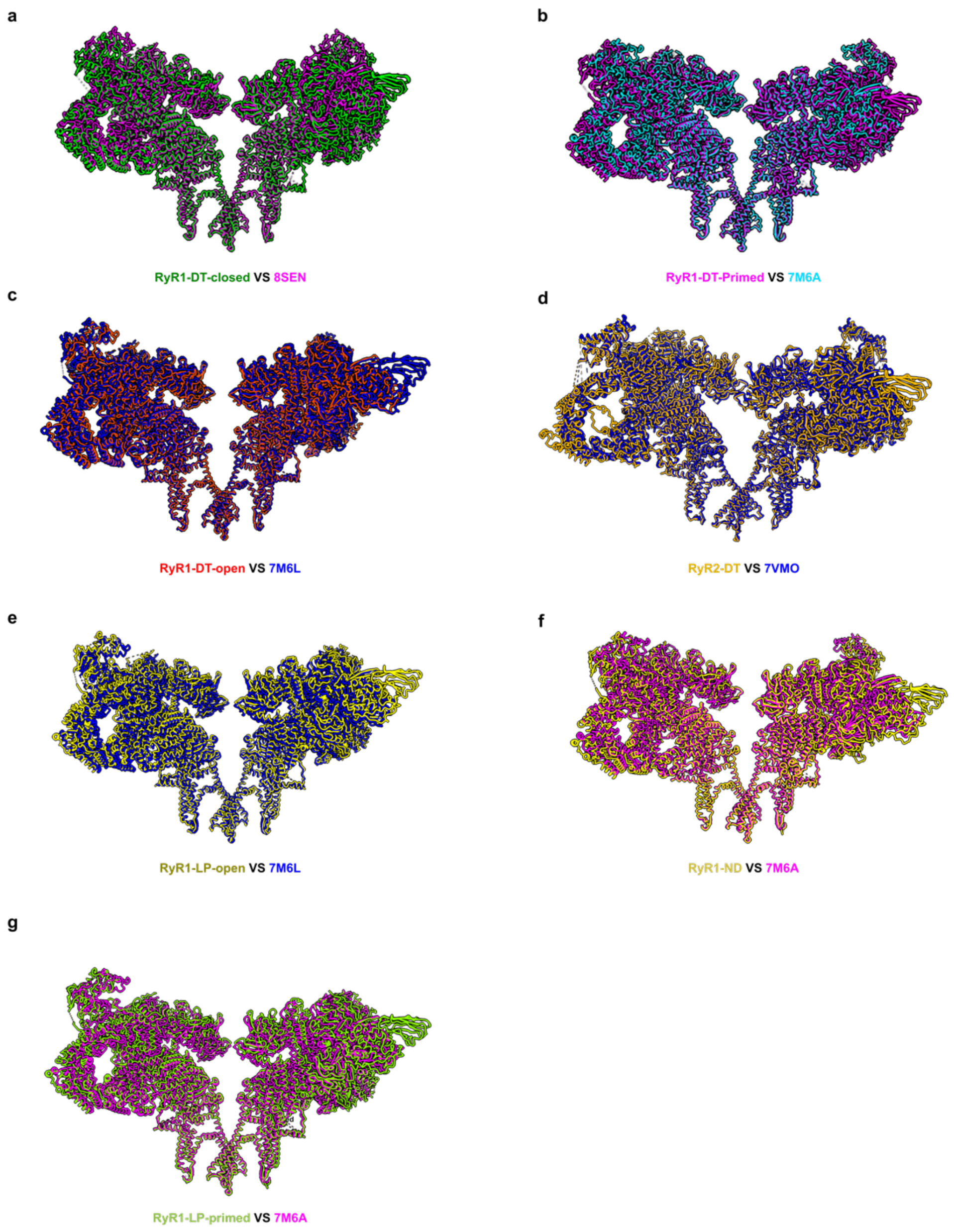


**Supplementary Figure 4 | Comparation of structures solved in this study with published structures of RyRs. a,** Model comparation of RyR1-DT-closed state to 8SEN, the overall RMSD is 4.5 Å. Pore region RMSD is 0.8 Å. **b,** Model comparation of RyR1-DT-primed state to 7M6A, the overall RMSD is 3.3 Å. Pore region RMSD is 0.7 Å. **c,** Model comparation of RyR2-DT state to 7VMO, the overall RMSD is 6.2 Å. Pore region RMSD is 0.6 Å. **e,** Model comparation of RyR1-LP-open state to 7M6L, the overall RMSD is 2.5 Å. Pore region RMSD is 1.5 Å. **f,** Model comparation of RyR1-ND state to 7M6L, the overall RMSD is 3 Å. Pore region RMSD is 0.8 Å. **g,** Model comparation of RyR1-LP-primed state to7M6A, the overall RMSD is 3.2 Å. Pore region RMSD is 1.3 Å.

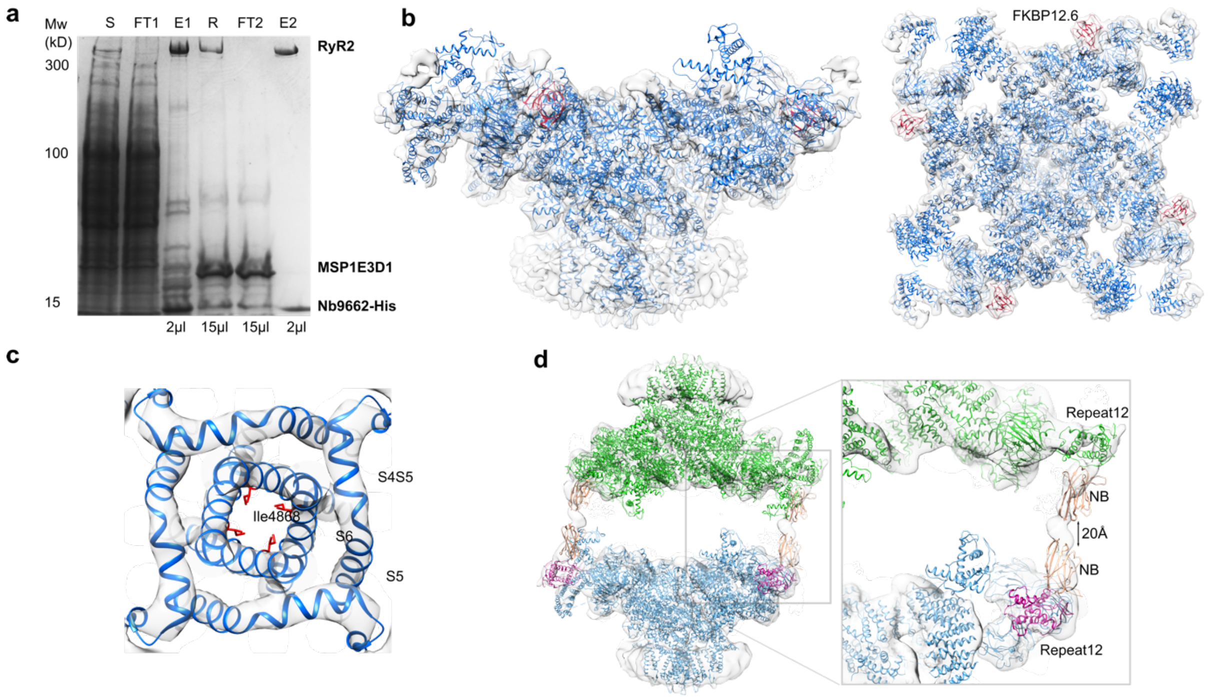


**Supplementary Figure 5 | Purification and low-resolution cryo-EM structure of bovine RyR2 a**, SDS-PAGE of sequential purification steps and reconstitution into lipid nanodiscs of bovine RyR2 Nb9662 complex. FT1- flow-through of the first purification step performed in detergent, E1- eluate in detergent, R- eluate 1 reconstituted into lipid nanodiscs, FT2- flow through of second purification step, E2- eluate of the second purification step. **b**, Cryo-EM reconstruction of purified bovine RyR2-FKBP12.6-Nb9662 complex at 7.5 Å resolution with a fitted structure of porcine RyR2-FKBP12.6 primed state (PDB: 6jH6, RyR2 in blue, FKBP12.6 in deep red). Side view and top views are shown. **c,** Top view of the gate region, the pore is in a primed state with an estimated distance of 12 Å between the C$\alpha$ atoms of gate residue Ile4868 in porcine RyR2. The gate residue Ile4868 (porcine RyR2) is shown in red. **d**, Side view of 3D reconstruction of a bovine RyR2 dimer at 13 Å resolution with a porcine RyR2 (PDB: 5GO9, blue/green) and Nb (PDB: 2X1O, beige) fitted into the map. The dimer interface is formed by the Nbs bound to the Repeat12 domain (magenta).

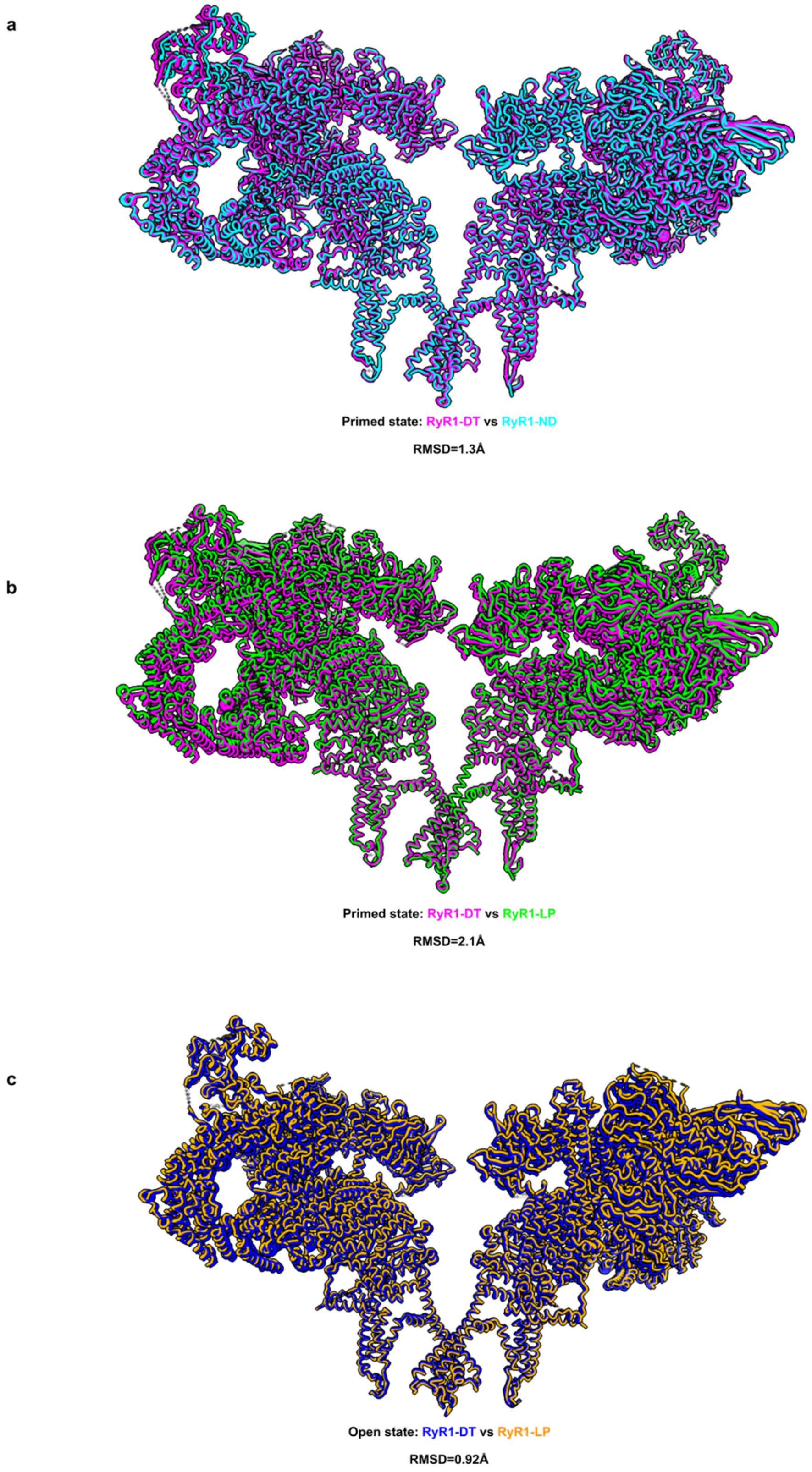


**Supplementary Figure 6 |Overall comparison of structures solved in this paper. a-b**, comparation of primed conformations solved in this study. **a**, RyR1-DT to RyR1-ND, **b**, RyR1-DT to RyR1-LP. **c**, Comparation of open conformation RyR1-DT to RyR1-LP.

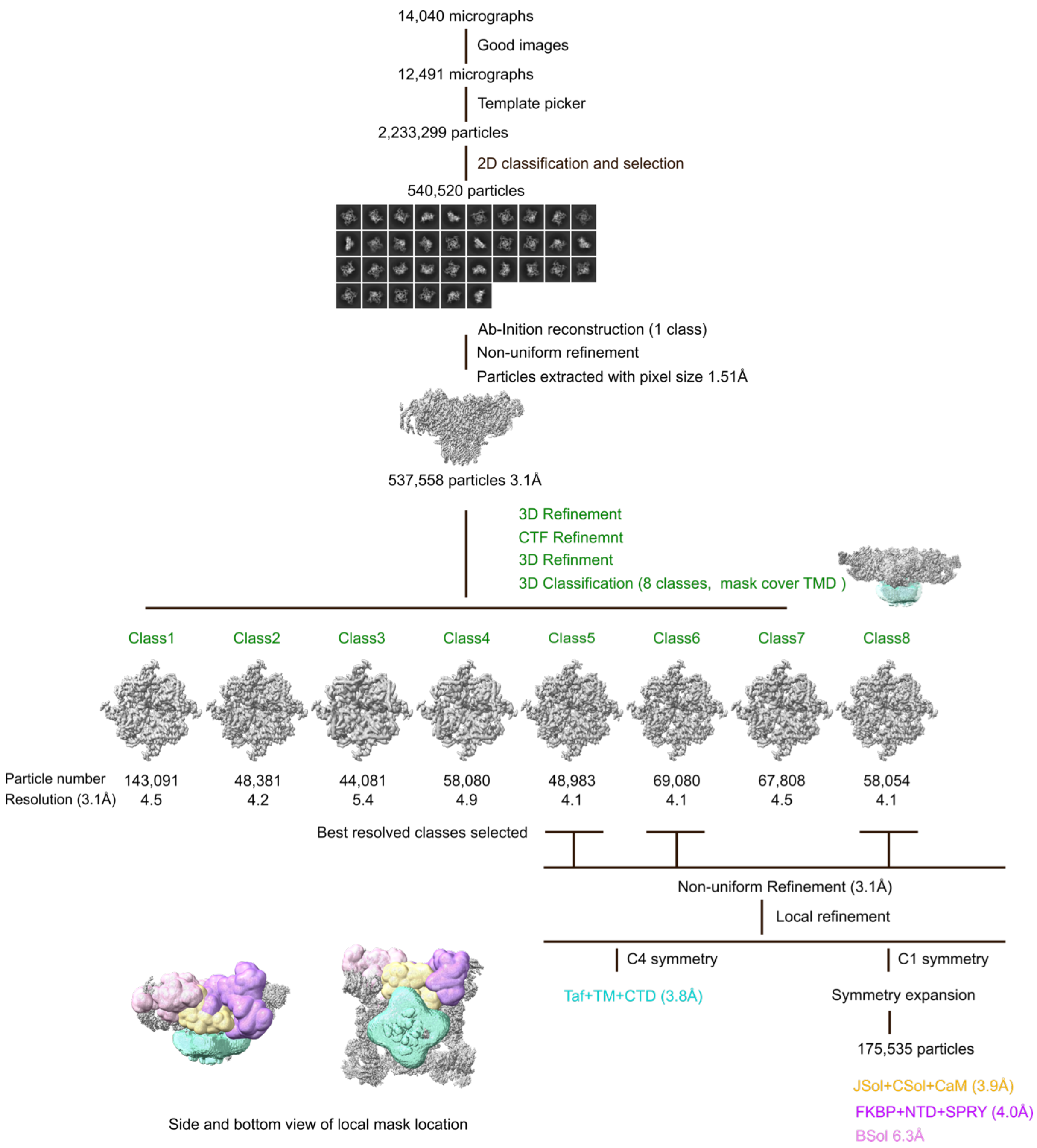


**Supplementary Figure 7 | Data processing scheme of the RyR-ND data set.** The green color indicates steps performed in Relion, other steps were performed in CryoSPARC. The masks used for 3D classification are indicated next to the 3D classification step. The masks used for local refinement are shown at the bottom.

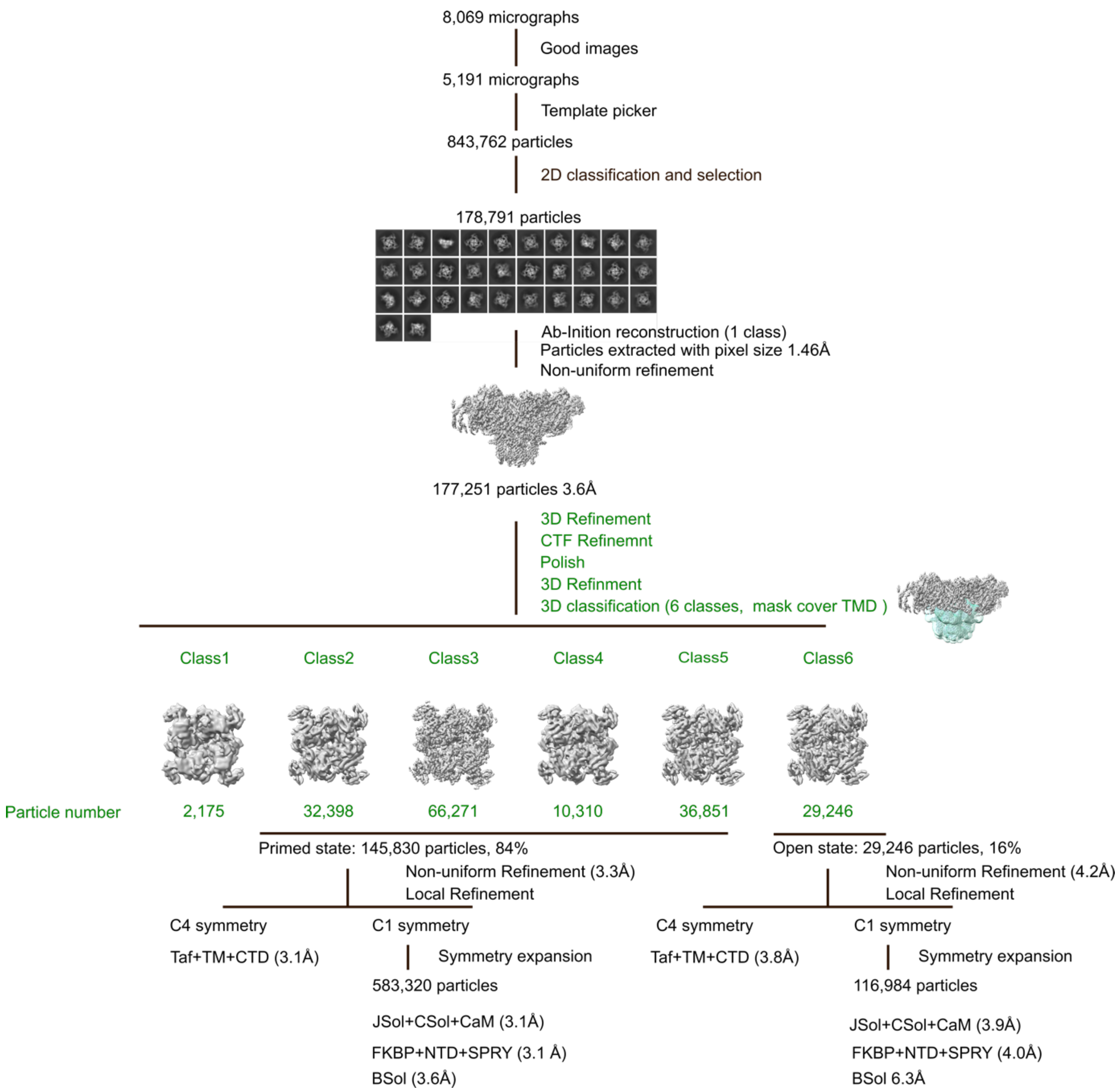


**Supplementary Figure 8 | Processing scheme for RyR1-DT with activators.** The green color indicates steps performed in Relion, other steps were performed in CryoSPARC. Masks used for local refinement are the same as used for RyR1-ND data processing.

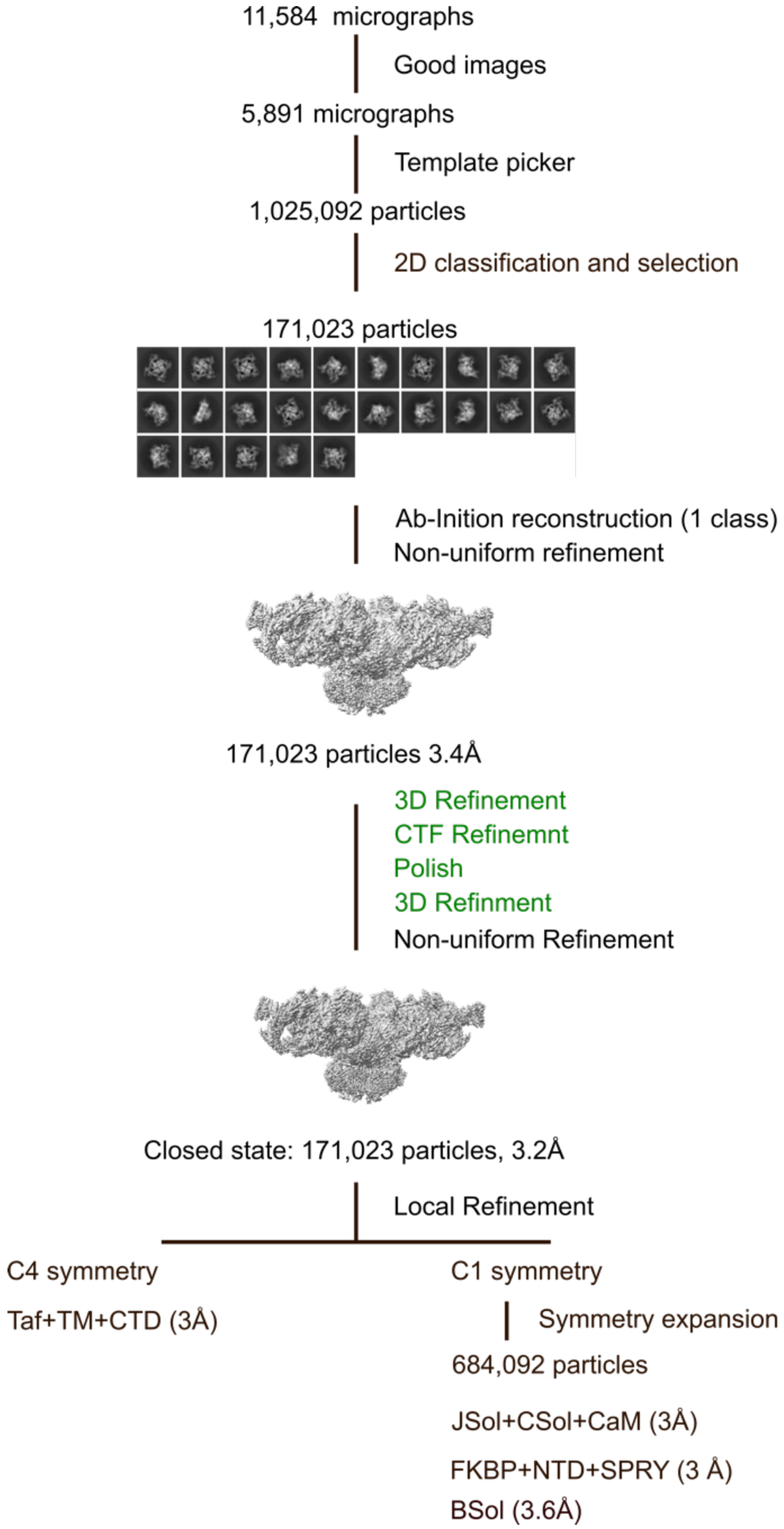


**Supplementary Figure 9 |Data processing of RyR1-DT data set without activators.** The green color indicates steps performed in Relion, other steps were performed in CryoSPARC. Masks used for local refinement are the same as in RyR1-ND data processing scheme.

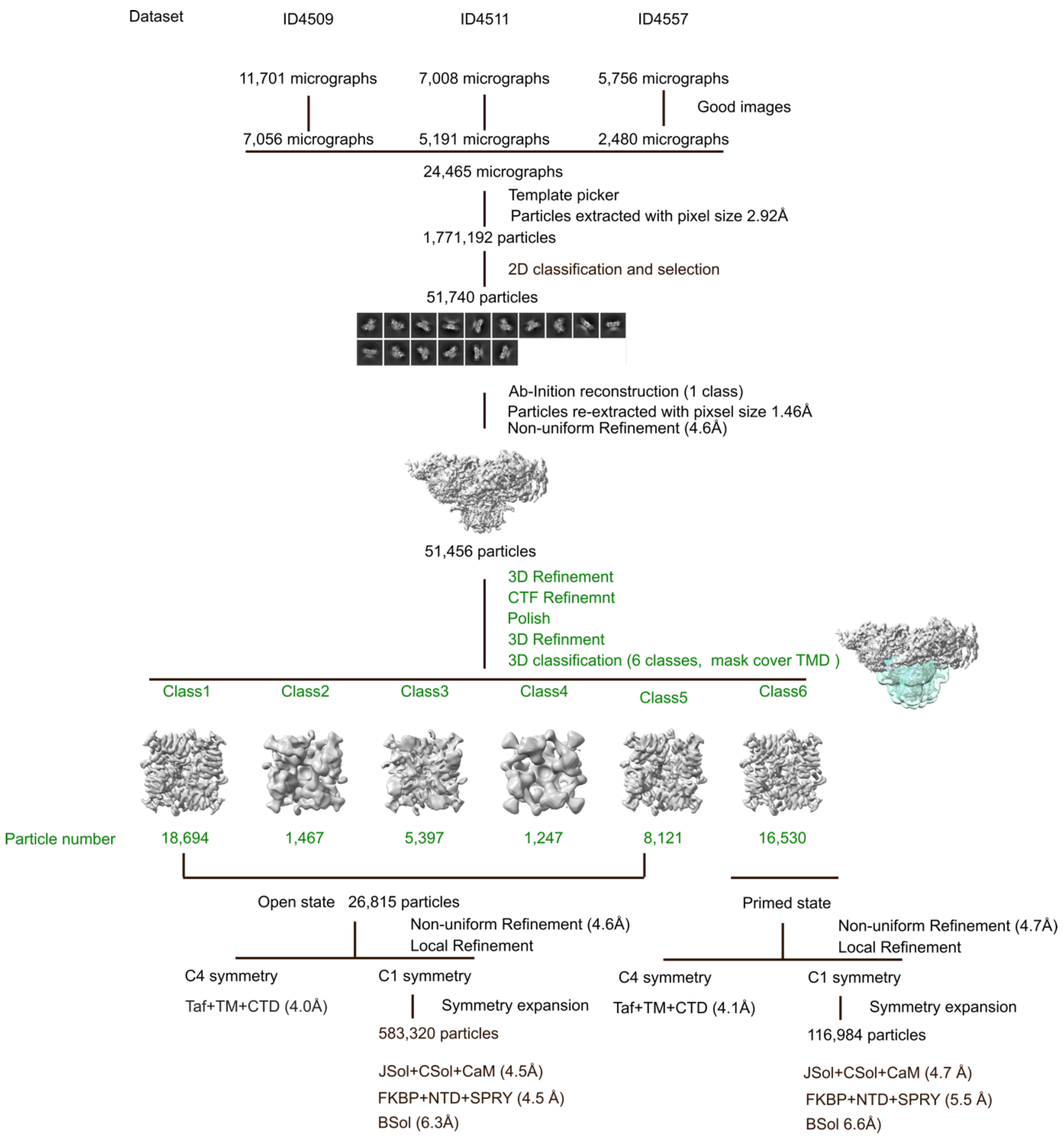


**Supplementary Figure 10 |Data processing of RyR1-LP data set**. The green color indicates steps performed in Relion, other steps were performed in CryoSPARC. Masks used for local refinement are the same as in RyR1-ND data processing scheme.

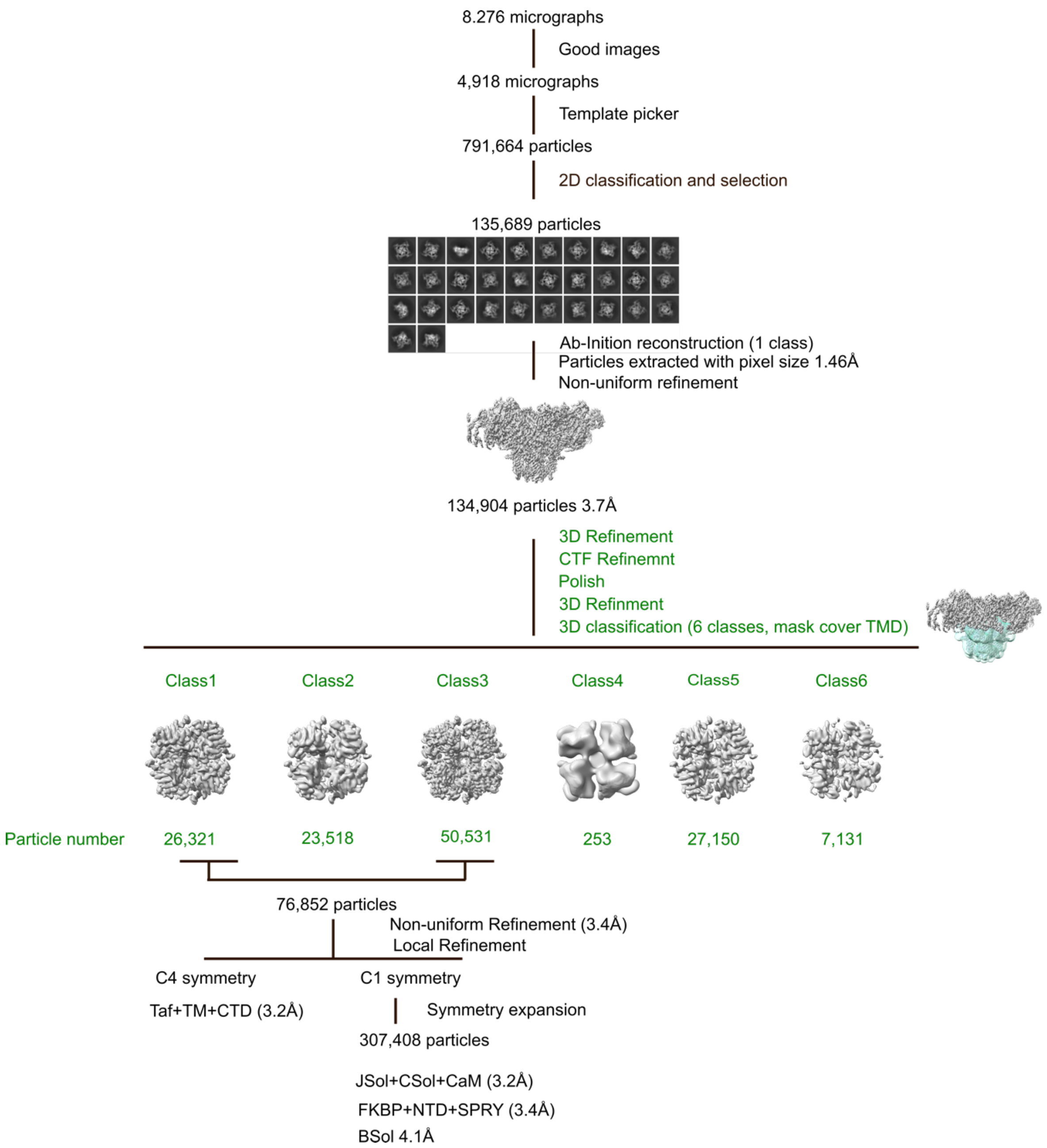


**Supplementary Figure 11 |Data processing of RyR2-DT data set.** The green color indicates steps performed in Relion, other steps were performed in CryoSPARC. Masks used for local refinement are the same as in RyR1-ND data processing scheme.

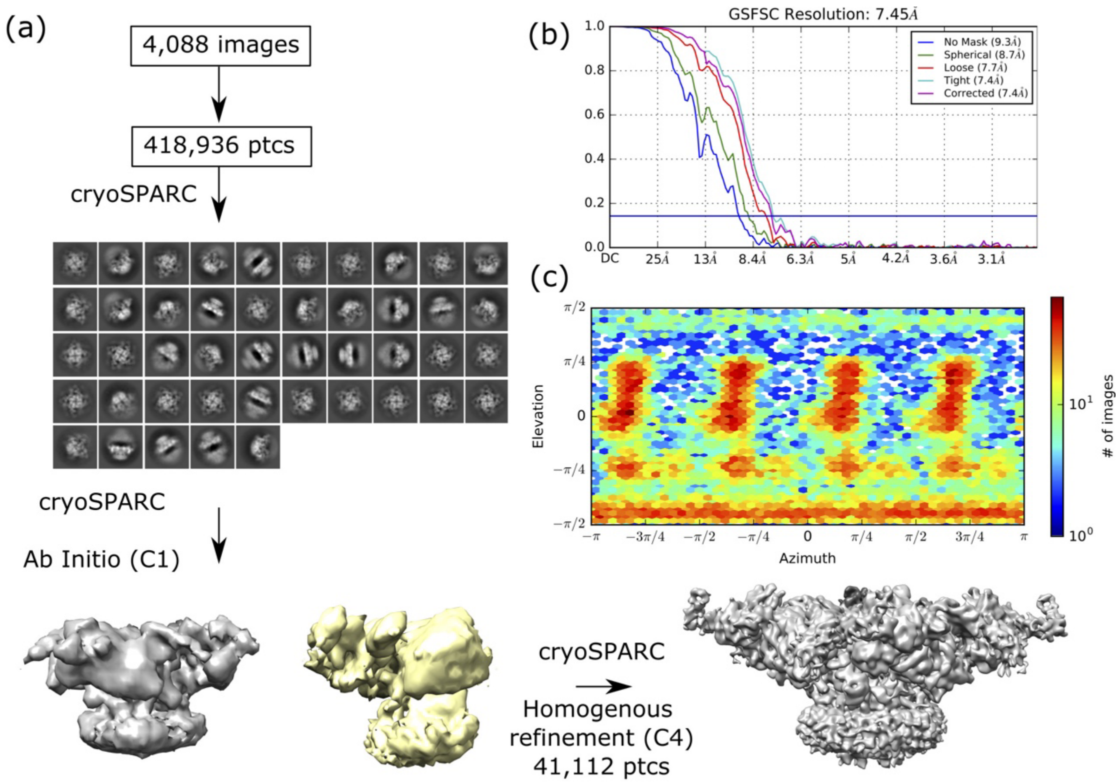


**Supplementary Figure 12 | Data processing of bovine RyR2-ND data set.** **a**, Scheme summarizing processing protocol used to obtain 3D reconstruction of RyR2 monomer. **b**, FSC curves of the final reconstruction. **c**, Angular distribution of particle orientations.

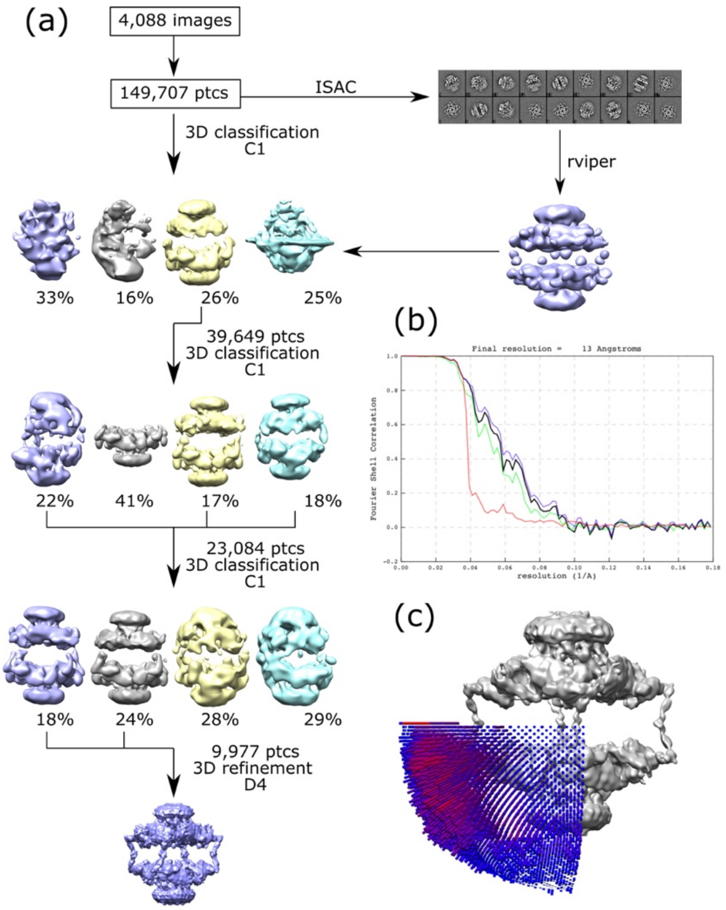


**Supplementary Figure 13 | Data processing of bovine dimeric bovine RyR2-ND data set.** **a**, Scheme summarizing processing protocol used to obtain 3D reconstruction of RyR2 dimers. **b**, FSC curves of the final reconstruction. **c**, Angular distribution of particles reconstructed applying D4 symmetry.
